# Phase transition specified by a binary code patterns the vertebrate eye cup

**DOI:** 10.1101/2021.08.12.455556

**Authors:** Revathi Balasubramanian, Xuanyu Min, Peter M.J. Quinn, Quentin Lo Giudice, Chenqi Tao, Karina Polanco, Neoklis Makrides, John Peregrin, Michael Bouaziz, Yingyu Mao, Qian Wang, Bruna L Costa, Diego Buenaventura, Fen Wang, Liang Ma, Stephen H Tsang, Pierre J. Fabre, Xin Zhang

## Abstract

The developing vertebrate eye cup is partitioned into the neural retina (NR), the retinal pigmented epithelium (RPE) and the ciliary margin (CM). By single cell analysis, we showed that a gradient of FGF signaling regulates demarcation and subdivision of the CM and controls its stem cell-like property of self-renewal, differentiation and survival. This regulation by FGF is balanced by an evolutionarily conserved Wnt signaling gradient induced by the lens ectoderm and the periocular mesenchyme, which specifies the CM and the distal RPE. These two morphogen gradients converge in the CM where FGF signaling promotes Wnt signaling by stabilizing β-catenin in a GSK3β-independent manner. We further showed that activation of Wnt signaling converts the NR to either the CM or the RPE depending on the level of FGF signaling. Conversely, activation of FGF transforms the RPE to the NR or CM dependent on Wnt activity. We demonstrated that the default fate of the eye cup is the NR, but synergistic FGF and Wnt signaling promotes CM formation both in vivo and in retinal organoid culture of human iPS cells. Our study reveals that the vertebrate eye develops through phase transition determined by a combinatorial code of FGF and Wnt signaling.

## INTRODUCTION

The vertebrate neural retina (NR) is insulated from extra-ocular tissue by the retinal pigmented epithelium (RPE) and circumscribed by the ciliary body (CB) and the iris in the periphery, the latter two structures control the intraocular pressure and the light intake, respectively (*1, 2*). These ocular structures arise in the embryonic eye cup from corresponding zones of the NR, RPE and ciliary margin (CM) (Fig. 1A). Similarly, the Drosophila eye is insulated optically by the subretinal pigment layer (SRP) at its base and the pigment rim on the side, which is derived from the pupal eye disc similar to the NR (*3*). Remarkably, the molecular mechanism specifying peripheral eye structure also appears to be conserved. In Drosophila, Wingless secreted by the eye-encasing head capsule generates a gradient of Wnt signaling required for PR formation (*3*). In mouse, Wnt signaling mediated by β-catenin is necessary for development of both the RPE and the CM (*4–7*). Interestingly, constitutively active β-catenin transforms the NR to the PR in Drosophila, but only to the CM in mouse (*3, 7, 8*). Why Wnt signaling in mouse fails to produce ectopic RPE in the retina is not known.

**Figure 1.**
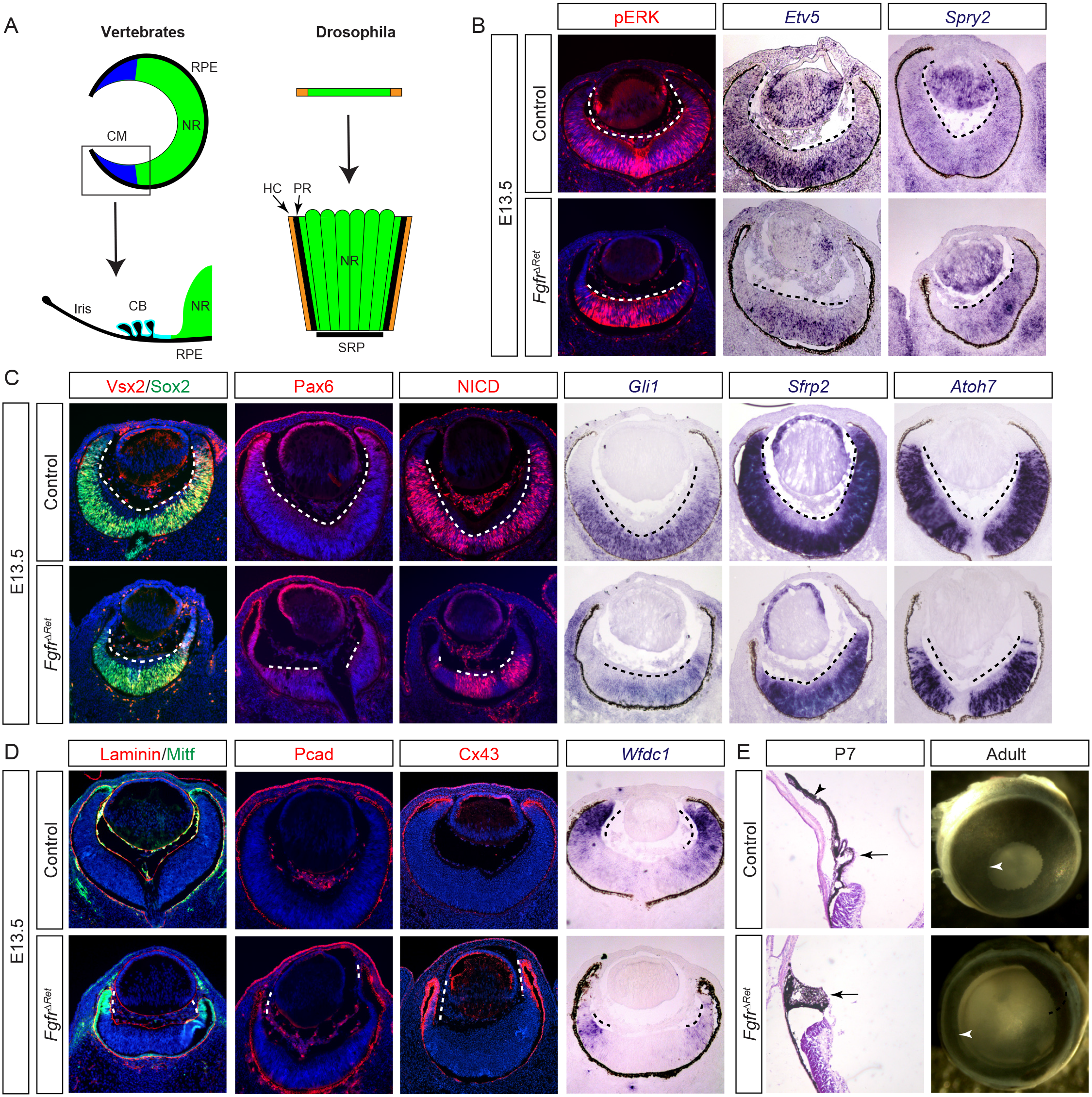
Genetic ablation of FGF receptors disrupted CM development. **(A)** The evolutionary conservation of peripheral ocular structures. The vertebrate eye cup is partitioned into the neural retina (NR), the retinal pigmented epithelium (RPE) and the ciliary margin (CM), the latter of which gives rise to the ciliary body (CB) and the iris. This resembles the Drosophila eye where the NR is shielded by the pigment rim (PR), the head capsule (HC) and the sub retinal pigment (SRP). **(B)** Deletion of FGF receptors in the peripheral retina abolished FGF signaling as indicated by the loss of pERK, *Etv5* and *Spry2* expression. **(C)** The NR domain (dotted lines) marked by Vsx2, Sox2, Pax6, N1CD, *Gli1, Sfrp2* and *Atoh7* expression was reduced in *Fgfr^ΔRet^* mutants. **(D)** *Fgfr^ΔRet^* mutant retina displayed ectopic expression of Mitf, Pcad and Cx43 (dotted lines), but down regulated *Wfdc1*, indicating loss of the CM domain. **(E)** *Fgfr^ΔRet^* mutant animals exhibited dysmorphic ciliary body (arrows) and iris hypoplasia (arrow heads) at P7 and aniridia (white arrowheads) in adulthood.

In vertebrate eye development, FGF is another transforming signal like Wnt. We and others have shown that genetic inactivation of FGF signaling disrupts formation of the NR and the optic disc (*9–11*). In contrast, ectopic expression of FGF not only transforms the RPE to the NR (*12–15*), but also induces markers of the CM in the junction between the ectopic NR and the remaining RPE (*16*). It is unclear, however, whether the ectopic CM is induced directly by FGF or indirectly by the ectopic NR. In fact, based on experiments performed in retinal organoids, Sasai and colleagues proposed that mutual inhibition of FGF and Wnt pivots the retinal progenitor cells to either the NR or RPE fate, leaving the NR-RPE boundary tissue to self-organize into the CM (*17*). In the current study, we show that, unlike in Drosophila, murine embryos have opposing morphogen gradients of FGF and Wnt in the peripheral eye cup induced by their distinctive sources of ligands in the NR and the lens ectoderm, respectively. Single-cell analysis further revealed that FGF signaling promotes subdivision of the CM zone in a dose dependent manner. Unexpectedly, FGF signaling is required to maintain the high level of Wnt activity necessary for CM fate. By genetic manipulation in mouse and chemical induction in retinal organoids, we demonstrate that the vertebrate eye cup is specified in a phase transition mode that can be programed reversibly by a binary code of FGF and Wnt signaling.

## RESULTS

### FGF signaling is required for CM development

During embryonic development, we observed a proximal^high^-distal^low^ pattern of pERK in E13.5 retinae (Fig. 1B), suggesting a gradient of FGF signaling (*18*). To examine its functional significance, we ablated FGF receptors 1 and 2 using *Pax6 α-Cre*, which is active in the peripheral retina beginning at E10.5 (*19*). In concordance with the role of FGF in inducing ERK signaling, both phosphorylation of ERK and expression of FGF-response genes *Etv5* and *Spry2* were lost in distal retinae of *α-Cre*;*Fgfr1^flox/flox^;Fgfr2^flox/flox^* (*Fgfr^ΔRet^*) mutants (Fig. 1B), leading to significant reduction in the retinal progenitor cell domain marked by Vsx2, Sox2, Pax6, Notch1 intracellular domain (NICD), *Gli1* and *Sfrp2* (Fig. 1C). As a result, the zone of the *Atoh7-*positive NR was also contracted, whereas the RPE markers Mitf, Pcad and Cx43 invaded deeply into *Fgfr^ΔRet^* mutant retinae, followed by a striking reduction in the pan-CM marker *Wfdc1* (Fig. 1D). At P7, control pups displayed well-organized ciliary bodies and elongated iris, but only clumps of pigmented cells were present at the tip of *Fgfr^ΔRet^* mutant retinae (Fig. 1E). In accordance with this, adult *Fgfr^ΔRet^* mice displayed almost complete aniridia (lacking the iris), demonstrating the critical role of FGF signaling in CM development.

### Single-cell analysis reveals that FGF signaling controls self-renewal, differentiation and survival of CM progenitors

To investigate the molecular basis of FGF signaling in regulating CM development, we performed single-cell RNA-sequencing (scRNAseq) of E13.5 eye cups. Taking advantage of an IRES-GFP cassette embedded in the *Pax6 α-Cre* driver, we used flow cytometry to enrich for Cre/GFP-expressing cells from peripheral retinae (Fig. 2A) and sequenced 6,628 control and 4,607 *Fgfr^ΔRet^* mutant cells at the mean depth of 2,811 genes per cell. As expected, unsupervised clustering analysis identified distinctive groups of retinal progenitor cells (RPC-1 to 4), neurogenic cells (Ngn-1, 2 and boundary) and differentiated retinal neurons (retinal ganglion cells (RGC), amacrine/horizontal cells (AC/HC) and photoreceptor cells (PRC), which were confirmed by expression of known molecular markers (Fig. 2B and S1A). Unlike previous scRNAseq analyses (*20, 21*), however, we were also able to discern three clusters that express CM specific markers *Mitf*, *Wls*, *Msx1* and *Wfdc1* (Fig. 2B and S1B). Projected onto the 2-dimensional UMAP plot, these CM cells formed a separate branch connected to the RPC clusters, diametrically opposite to the branch of retinal neurons (Fig. 2C). We next examined the trajectory of cell differentiation by RNA velocity analysis, which exploits dynamics of mRNA splicing to uncover the rate and direction of transcriptomic changes (*22*). It correctly identified the differentiation path from the RPC to either the NR or the CM fates (Fig. 2D), which correlated with the transitioning of the cell cycle from the S phase via the G2M to the G1 phase (Fig. 2E). Interestingly, our analysis showed that once progenitor cells on the NR trajectory entered the neurogenic state in the G2M phase, they irreversibly progressed toward terminal differentiation. On the CM trajectory, however, progenitor cells (RPC-3) residing in the G2M phase could either move toward the terminal CM state or reenter the cell cycle to replenish the RPC pool. Thus CM progenitor cells displayed the hallmark of stem cells in their capacity to either self-renew or differentiate. In *Fgfr^ΔRet^* mutants, however, RPCs were biased against self-renewal in favor of the terminal CM fate. This was confirmed by comparing transition probabilities between control and mutant progenitor cells at similar locations within RPC clusters (Fig. 2F). Consistent with this, cell proliferation was significantly reduced in *Fgfr^ΔRet^* mutants (Fig. S2A) and the overall RNA velocity slowed down considerably in the circling RPC pool, but it accelerated in proximal and medial CM clusters before ramping down in the distal CM (Fig. 2G). This accounted for the increase in the percentage of CM cells at the expense of RPC cells (Fig. S2B). Therefore, FGF signaling controls the decision of self-renewal versus differentiation of CM progenitors.

**Figure 2.**
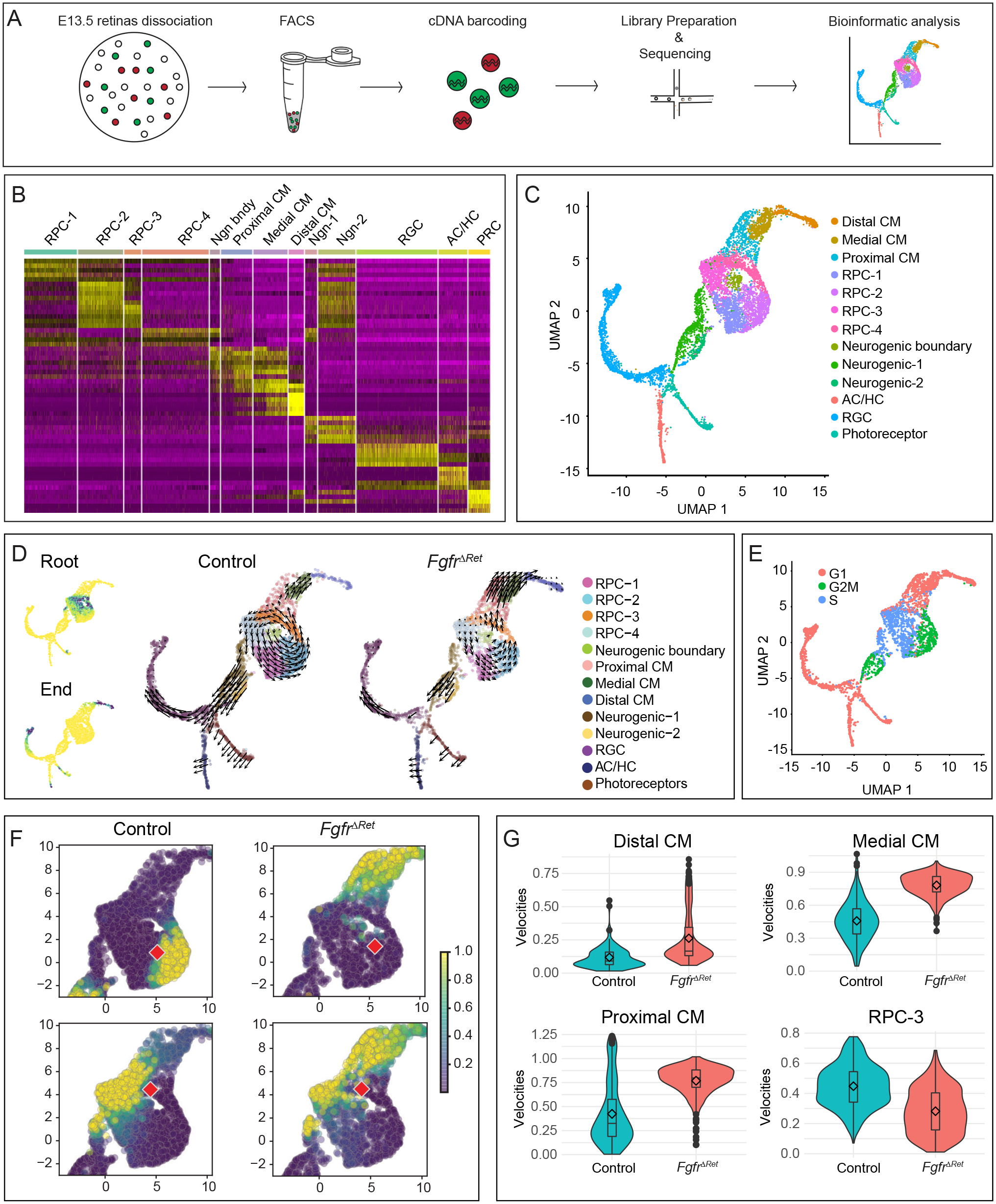
Single cell analysis showed that FGF signaling regulates self-renewal and differentiation of CM progenitors. **(A)** Schematic diagram of single-cell RNAseq. **(B)** Heat map of differential gene expression in single-cell clusters. RPC: retinal progenitor cell, RGC: retinal ganglion cell, AC: Amacrine cell, HC: Horizontal cell. PR: photoreceptor. **(C)** UMAP representation of single-cell clusters. **(D)** Left: Diffusion analysis of RNA velocities identified the root and the end of cell differentiation as represented by high density regions (dark green) after forward and reverse Markov processes. Right: Cell differentiation trajectories were revealed by the velocity field projected on the UMAP plot. Arrows indicate local RNA velocities on a regular grid. **(E)** The state of the cell cycle represented on the UMAP plot. **(F)** Single-step transition probabilities from the starting cells (red square) to neighboring cells showed the bias of CM progenitors in *Fgfr^ΔRet^* mutants toward differentiation against self-renewal. **(G)** Quantification of single-cell velocities in each cluster showed the accelerated differentiation and reduced proliferation of CM progenitor cells.

We next asked whether FGF signaling is also required for terminal differentiation of the CM. Our scRNAseq analysis classified CM cells into three clusters: the distal CM defined by expression of *Wls* and *Otx2*, the medial CM by *Msx1* and the proximal CM by overlapping expression of *Sox2* and *Cdo* (Fig. 3A). We showed that this in silico segregation of CM cells corresponds to their spatial separation in situ (*23*), as expression of Wls and Otx2 extended beyond the Pcad-positive RPE into the distal tip of the retina, bordering the domain of Msx1 expression (Fig. 3B). Msx1 in turn overlapped with Sox2 and Cdo, the latter two reaching further into the center of the retina. In contrast, *Fgfr^ΔRet^* mutant retinae displayed encroachment of Pcad, expansion of Wls and Otx2, loss of Msx1 and retreat of Sox2 and Cdo. This was in agreement with scRNAseq analysis of *Fgfr^ΔRet^* mutants that *Wls* and *Otx2* expanded from the distal CM into the medial or even proximal CM domains at the expense of *Msx1*, *Sox2* and *Cdo*. These results demonstrated that FGF signaling is required for spatial demarcation and molecular distinction of the CM subdivisions.

**Figure 3.**
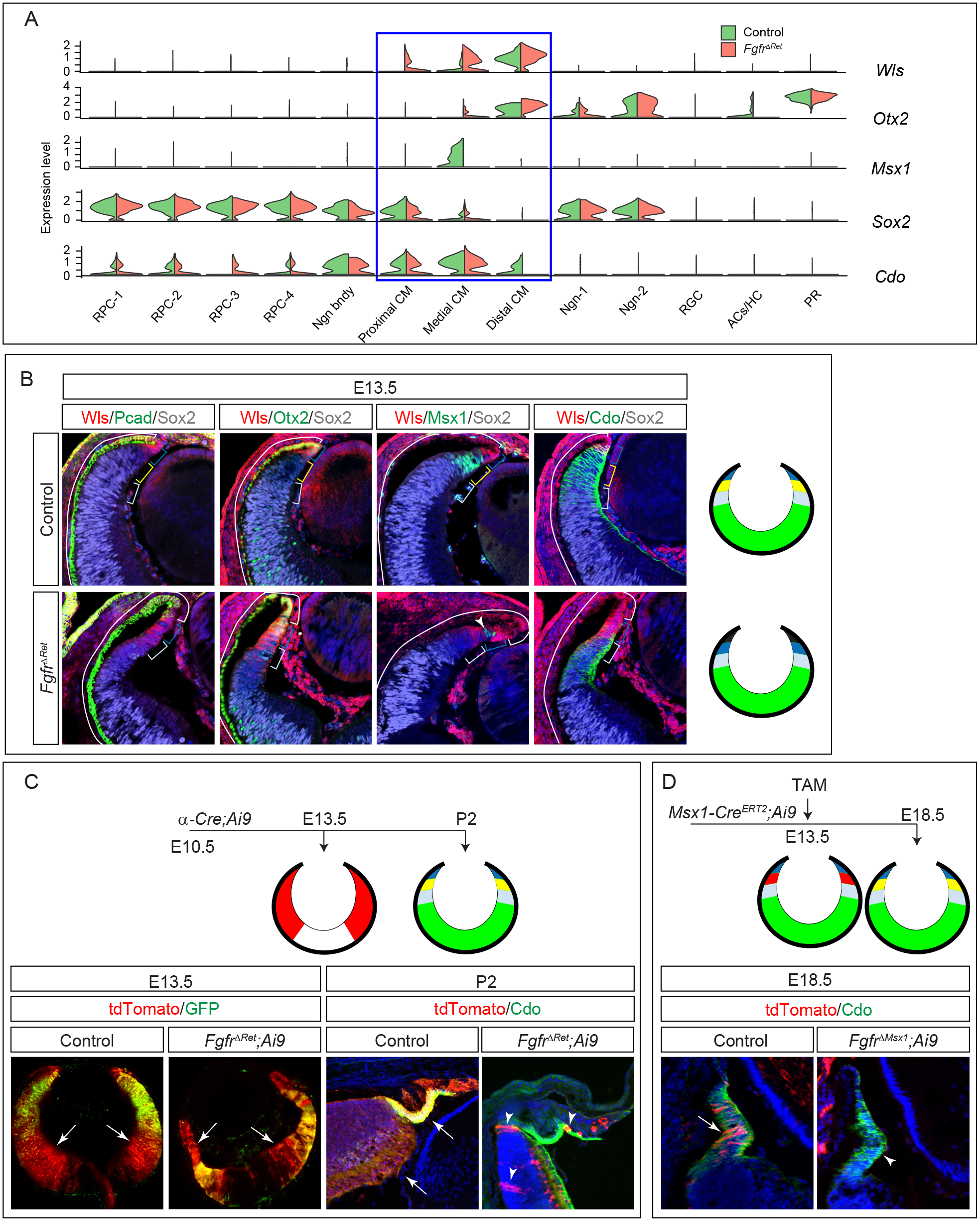
FGF signaling is required for CM subdivision and survival. **(A)** Violin plots showed that the CM can be subdivided by the overlapping expression of *Wls*, *Otx2*, *Msx1*, *Sox2* and *Cdo*, all of which were dysregulated in *Fgfr^ΔRet^* mutants. **(B)** Immunostaining confirmed that the in silico clustering of CM cells matches the spatial separation of CM subdomains. Aberrant invasion of Pcad, expansion of Wls and Otx2, loss of Msx1 and reduction in Sox2 and Cdo in *Fgfr^ΔRet^* mutants demonstrated CM differentiation defects. Brackets indicate the domain of the RPE and the distal, medial and proximal CM, corresponding to black, dark blue, yellow and light blue regions in diagrams on the right. The NR is indicated in green. Arrowhead marks residual wild type cells still expressing Msx1. **(C)** At E13.5, although *Pax6 α-Cre* was expressed only in the peripheral retina indicated by its GFP reporter, it has already activated tdTomato expression (arrows) from the *Ai9* Cre reporter throughout the retina. At P2, these tdTomato positive progenies remained in control retinae, but only a few (arrowheads) were left in *Fgfr^ΔRet^* mutants. The tdTomato expression is indicated in red in diagrams. **(D)** CM cells were pulse labeled by tamoxifen induction of *Msx1-Cre^ERT2^* at E13.5 and detected at E18.5 by tdTomato expression from the *Ai9* Cre reporter. Although these cells remained in the control CM identified by Cdo expression (arrow), they had largely disappeared in the *Fgfr^ΔRet^* mutant (arrowhead), suggesting cell survival defects.

Lastly, we examined the potential role of FGF signaling in retinal cell survival. To this end, we permanently labeled *α-Cre*-expressing cells at E10.5 using the *Ai9* Cre reporter and followed their cell fates through development by the tdTomato expression (Fig. 3C). At E13.5, we observed tdTomato positive cells spread to much of the retina in both control and *Fgfr^ΔRet^* mutants, even beyond the *α-Cre* expressing domain marked by GFP expression. Although control retinae at P2 still displayed ubiquitous expression of tdTomato, only a handful of tdTomato+ cells persisted in *Fgfr^ΔRet^* mutants, likely a result of incomplete recombination due to Cre mosaicism. To determine whether loss of tdTomato+ cells in *Fgfr^ΔRet^* mutants was solely due to depletion of the RPCs, we selectively labeled medial CM cells by inducing *Msx1-Cre^ERT2^* with Tamoxifen at E13.5 and examined their fates at E18.5 (Fig. 3D). In contrast to abundant expression of tdTomato reporter in control ciliary bodies marked by Cdo, *Msx1-Cre^ER2^*;*Fgfr1^flox/flox^;Fgfr2^flox/flox^;Ai9* (*Fgfr^ΔMsx1^*;*Ai9*) mutants were largely devoid of tdTomato+ cells. Therefore, FGF signaling is required for survival of CM cells even after their onset of differentiation.

### The ciliary margin is patterned by graded FGF signaling

We next focused on how FGF signaling regulates patterning of the CM. Our scRNAseq analysis revealed that the retina expresses three FGF ligands - *Fgf3* restricted to the RPCs, *Fgf9* extending into the proximal CM and *Fgf15* further encompassing the medial CM (Fig. 4A). Intriguingly, this nested pattern of *Fgf* expression coincides with the proximal-distal gradient of FGF signaling (Fig. 1B) and subdivision of the CM in the distal retina. We hypothesized that sequential removal of Fgfs may flatten the gradient of FGF signaling and disrupt the patterning of the CM (Fig. 4B). To test this model, we first deleted *Fgf9*, which was previously shown to control the anterior boundary of the RPE (*24*). In *α-Cre*;*Fgf9^flox/flox^* (*Fgf9^ΔRet^*) mutants, *Mitf* expression advanced from the distal CM toward the central retina, accompanied by the down regulation of *Vsx2*, *Wfdc1* and *Msx1* expression in the proximal CM domains. We next ablated both *Fgf3* and *Fgf9* in *α-Cre*;*Fgf3^flox/flox^*;*Fgf9^flox/flox^* (*Fgf3/9^ΔRet^*) mutants, which led to further encroachment of *Mitf* into the retina accompanied by a reduction in *Vsx2* and *Wfdc1* and loss of the medial CM maker *Msx1*. These results suggested that the graded expression of *Fgfs* in the distal retina is necessary for the subdivision of the CM.

**Figure 4.**
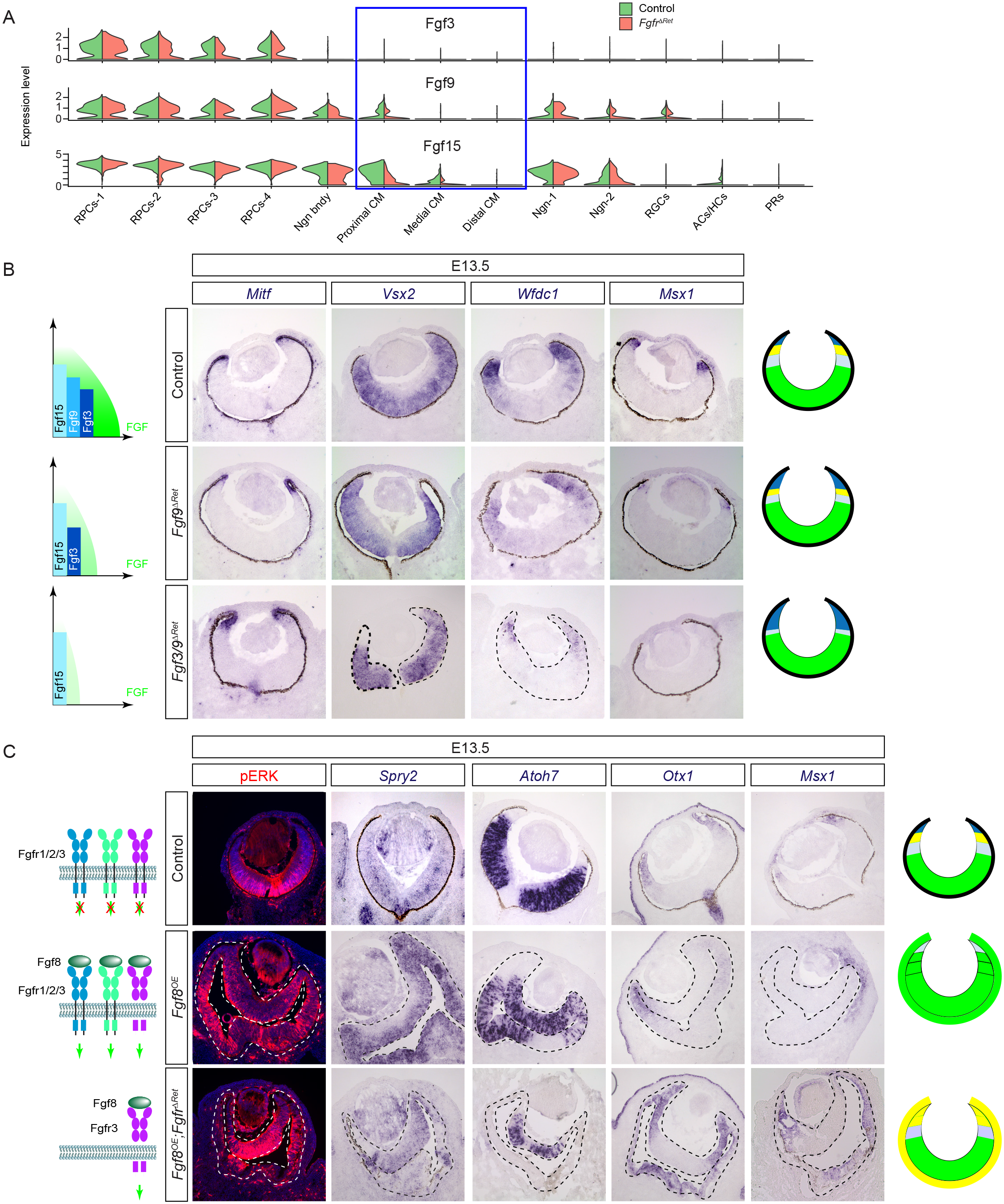
CM development is dependent on the dosage of FGF signaling. **(A)** scRNAseq analysis revealed a nest pattern of *Fgf3*, *Fgf9* and *Fgf15* expression in the retina. **(B)** By reducing the dosage of Fgf ligands, *Fgf9^ΔRet^* and *Fgf3/9^ΔRet^* exhibited progressive expansion of the distal CM/RPE marker *Mitf*, reduction in the proximal CM/NR marker *Vsx2*, and loss of pan CM marker *Wfdc1* and medial CM marker *Msx1*. **(C)** Overexpression of Fgf8 in *Fgf8^OE^* mutants stimulated pERK and FGF-responsive gene *Spry2* in the presumptive RPE territory, which was transformed to NR as indicated by *Atoh7* expression. Reducing the strength of FGF signaling by deleting Fgfr1 and 2 in *Fgfr^ΔRet^;Fgf8^OE^* mutants down regulated pERK. This prevented expression of *Atoh7*, but induced the CM markers *Otx1* and *Msx1*, indicating the transformation to the CM fate.

Previous studies have shown that over activation of FGF signaling can transform the RPE into the NR. Since our above result indicated that the dosage of *Fgfs* is important for the specification of the CM in the retina, we wondered if titration of FGF signaling may redirect the RPE to the CM fate. To this end, we first crossed *α-Cre* with *R26^LSL-Fgf8^* to induce ectopic expression of Fgf8 in the retina, which is expected to diffuse to the RPE to activate FGF receptors (Fgfr1-4). This was confirmed by the ectopic induction of pERK and FGF response gene *Spry2* throughout the RPE/NR double-layered eye cup in *α-Cre*;*R26^LSL-Fgf8^* (*Fgf8^OE^*) embryos (Fig. 4C). Furthermore, the original RPE layer now resembled the NR in both the thickness and the *Vsx2^+^Atoh7^+^Mitf ^-^Gja1^-^* expression pattern, confirming its conversion into the NR. We next attenuated the ectopic FGF signaling in the former RPE territory by ablating Fgfr1 and 2, leaving only Fgfr3 and 4 available to transmit the Fgf8 signal. Indeed, this genetic manipulation reduced but did not eliminate pERK and *Spry2* staining in the presumptive RPE in the *α-Cre*; *Fgfr1^flox/flox^;Fgfr2^flox/flox^;R26^LSL-Fgf8^* (*Fgfr^ΔRet^;Fgf8^OE^*) embryo. Although this region remained thickened compared to the control RPE, it no longer expressed the NR marker *Atoh7*, but instead acquired CM markers *Otx1* and *Msx1*. Taken together, these results showed that graded FGF signaling is important for CM differentiation in both the RPE and the retinal domains.

### FGF signaling determines the transformative activity of Wnt signaling

Since the above results demonstrate an essential role of FGF signaling in CM development, we wondered whether FGF interacts with Wnt signaling, which is also known to promote the CM fate (*7*). Analyzing our scRNAseq data, we noticed that *Lef1* and *Axin2*, two response genes for canonical Wnt signaling, were down regulated in *Fgfr^ΔRet^* mutants, suggesting that FGF signaling may be required for Wnt activity in the peripheral retina (Fig. 5A). This was unexpected because previous studies have suggested that FGF and Wnt play opposing roles in early eye cup patterning (*6, 17*). We thus sought to confirm our findings by mosaic analysis. As described above, inactivation of FGF signaling resulted in expansion of Mitf and Otx2 expression into the *Fgfr^ΔRet^* peripheral retina (Fig. 5B). Due to the mosaic activity of *α-Cre*, however, we occasionally observed patches of cells still negative for Mitf and Otx2 in distal retinae, suggesting that they were residual wild type cells. Remarkably, Lef1 expression was retained in these Mitf^-^/Otx2^-^ cells but lost in their neighbors (Fig. 5B, arrowheads and dotted line), demonstrating that FGF signaling cell-autonomously regulates Wnt signaling.

**Figure 5.**
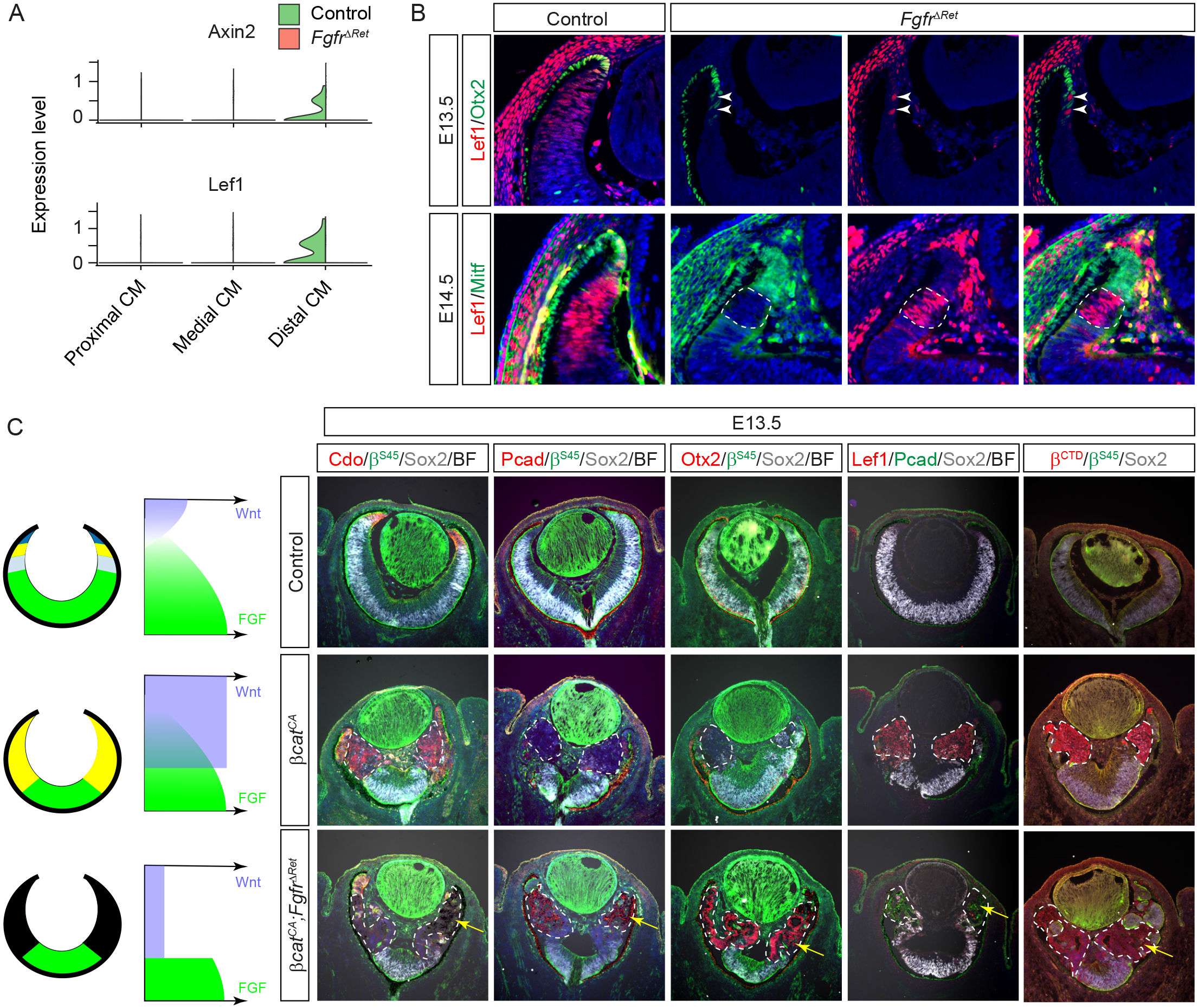
FGF signaling determines the CM fate by promoting Wnt signaling. **(A)** Violin plots showed down regulation of Wnt response genes *Lef1* and *Axin2* in the *Fgfr^ΔRet^* mutant transcriptome. **(B)** In mosaic analysis, *Fgfr^ΔRet^* mutant cells acquired ectopic expression of Otx2 and Mitf at the expense of Lef1. In contrast, the remaining wild type cells identified by the lack of Otx2 and Mitf expression still maintained Lef1 expression. **(C)** Constitutively activation of Wnt signaling by deleting the β-catenin^Ser45^ (β^Ser45^) motif transformed *βcat^CA^* retinae to the CM as indicated by Cdo expression. Further deletion of FGF receptors in *Fgfr^ΔRet^*;*βcat^CA^* mutant retinae converts them to the RPE as indicated by ectopic expression of Pcad and Otx2 as well as the appearance of pigmentation (arrows). Both the level of Lef1 and the amount of β-catenin detected by the β-catenin C-terminal antibody (β^CTD^) arose in *βcat^CA^* retinae but declined in *Fgfr^ΔRet^*;*βcat^CA^* mutants, showing that the Wnt-β-catenin signaling is dependent on FGF signaling. The mutant regions are marked by dotted lines.

We next asked whether FGF signaling is also required functionally for Wnt signaling to promote CM formation. To test this hypothesis, we first induced constitutive Wnt signaling in the retina by deleting exon 3 of *β-catenin*, which encodes the Ser45 phosphorylation site necessary for GSK3β-regulated degradation (*25*). Identified by an antibody specific to the β-catenin^Ser45^ (β^Ser45^) residue, *β-catenin* mutant cells in *α-Cre*;*β-catenin^fl3/fl3^* (*βcat^CA^*) retinae up regulated the expression of the CM marker Cdo at the expense of Sox2 (Fig. 5C). This is consistent with previous reports that activation of Wnt signaling transforms the NR to the CM (*7, 8*). The Cdo expression, however, was abolished after the inactivation of FGF signaling in *α-Cre*; *Fgfr1^flox/flox^;Fgfr2^flox/flox^*;*β-catenin^fl3/fl3^* (*Fgfr^ΔRet^*;*βcat^CA^*) retinae. Instead, these *Fgfr^ΔRet^*;*β-catenin^CA^* cells acquired expression of Otx2 and Pcad as well as RPE-like pigmentation (Fig. 5C, arrows), confirming that they have taken on an RPE identity. Taken together, these results demonstrated that the level of FGF signaling dictates whether Wnt signaling transforms the NR to either the CM or the RPE fate.

To understand the mechanism by which FGF signaling regulates Wnt activity, we examined Lef1 expression induced by the stabilized β-catenin. As expected, Lef1 expression was greatly elevated in *βcat^CA^* retinae, but it became significantly attenuated after the loss of FGF signaling in *Fgfr^ΔRet^*; *βcat^CA^* retinae (Fig. 5C). This is consistent with our above finding that FGF signaling influences Lef1 expression in wild type retina (Fig. 5A and B). Moreover, we observed that *βcat^CA^* mutants exhibited strong cytoplasmic β-catenin staining as revealed by a β-catenin C-terminal antibody (β^CTD^), but this staining was markedly reduced in *Fgfr^ΔRet^*; *βcat^CA^* mutants. Since *βcat^CA^* mutants expressed the truncated β-catenin resistant to GSK3β phosphorylation, this result suggested that FGF signaling promotes the stability of β-catenin in an GSK3β-independent manner. To determine the downstream effector that mediates FGF signaling to promote Wnt activity, we further disrupted MAPK pathway by deleting *Mek1* and *2* in *α-Cre*;*Mek1^flox/flox^;Mek2^-/-^* (*Mek^ΔRet^*) mutants, which was confirmed by loss of pERK in the peripheral retina. Importantly, *Mek^ΔRet^* animals phenocopied *Fgfr^ΔRet^* mutants in their aberrant expression of *Vsx2*, *Atoh7*, *Wfdc1*, *Msx1* and *Mitf* during the embryogenesis and dysgenesis of the ciliary body and iris in adults (Fig. S3). Altogether, these results show that FGF-MAPK signaling stabilizes β-catenin to promote Wnt signaling in CM development.

### The lens ectoderm and periocular mesenchyme induce a Wnt signaling gradient to pattern the eye cup

In Drosophila development, the head capsule patterns the peripheral eye structures by secreting Wnt ligand to induce the Wnt signaling gradient (*3*). We wondered whether this mechanism of Wnt signaling induction is conserved in vertebrates. The vertebrate RPE consists of the proximal and distal domains (*2*), the latter forms the pigmented epithelium of the ciliary body lining the non-pigmented epithelium derived from the CM. Consistent with previous reports (*7, 26*), we found that Wnt signaling revealed by a highly sensitive TCF-GFP reporter is strongest in the distal CM and distal RPE before tapering off toward the center of the eye cup (Fig. 6A) (*27*). We reasoned that there could be three potential sources of Wnt - the distal retina, the periocular mesenchyme and the lens ectoderm (the surface ectoderm and the lens). These tissues can be targeted by *α-Cre*, *Wnt1-Cr*e and *Le-Cre*, respectively (Fig. 6A), allowing us to eliminate their Wnt production by genetically ablating Wntless, a cytoplasmic transporter necessary for Wnt secretion (*19, 26, 28, 29*). In both *α-Cre*;*Wntless^flox/flox^* (*Wls^ΔRet^*) and *Wnt1-Cr*e;*Wntless^flox/flox^* (*Wls^ΔPM^*) embryos, we did not observe any ocular defects (Fig. S4). In line with a previous report, however, *Le-Cre*;*Wntless^flox/flox^* (*Wls^ΔLE^*) mutants exhibited posteriorly shift of the RPE-retina boundary (Fig. 6B) (*26*). In addition, we observed that *Wls^ΔLE^* mutants lost the distal RPE marker *Wnt2b*, but not the proximal marker *Ttr*, suggesting a role of lens ectoderm-derived Wnt ligands in RPE regionalization (*30*). We further generated *Le-Cre*;*R26^LSL-Wnt1^* to overexpress *Wnt1* in the lens ectoderm, which abolished lens development due to its inhibitory effect in lens induction (Fig. 6C) (*31*). In the eye cup, however, this led to the up regulation of *Wnt2b* without affecting *Ttr* expression. Moreover, the distal eye cup in *Wls^ΔLE^* mutants expressed RPC marker *Vsx2*, but none of the pan CM markers *Wfdc1* and *Otx1*, the medial CM marker *Msx1* and distal CM marker *Mitf* (Fig. 6D). Taken together, these results showed that the lens ectoderm-derived Wnt is responsible for specification of both the distal RPE and the CM.

**Figure 6.**
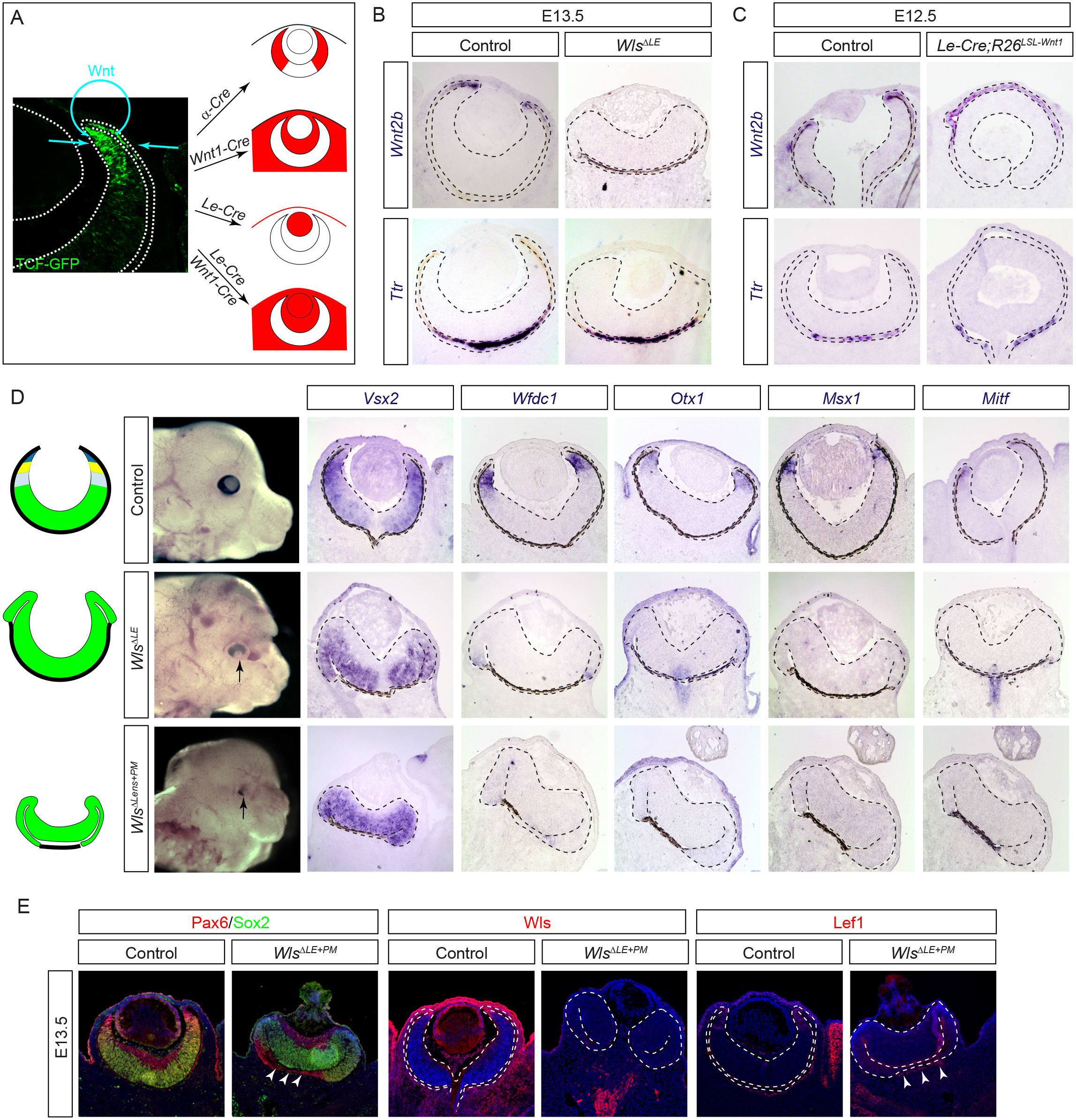
Paracrine Wnt signaling patterns the CM and the distal RPE. **(A)** The potential source of Wnt signaling gradient revealed by the TCF-GFP reporter may be targeted by *α-Cre* for retina, *Wnt1-Cr*e for periocular mesenchyme and *Le-Cre* for the lens ectoderm (the surface epithelium and the lens). **(B)** Ablation of Wnt transporter Wls in the lens ectoderm specifically disrupted the proximal RPE differentiation in *Wls^ΔLE^* mutants as indicated by loss of *Wnt2b*, but not the proximal RPE marker *Ttr*. **(C)** Overexpression of *Wnt1* in the lens ectoderm also affected the proximal RPE by expanding *Wnt2b* expression in *Le-Cre*;*R26^LSL-Wnt1^* eye cup without affecting *Ttr* expression. **(D)** The CM was lost in *Wls^ΔLE^* mutants as shown by the expansion of *Vsx2* and absence of *Wfdc1*, *Otx1*, *Msx1* and *Mitf*. The band of the RPE was further diminished in *Wls^ΔLE+ΔPM^* mutants. **(E)** *Wls^ΔLE+ΔPM^* mutant eyes lost Wls and Lef1 in both the lens ectoderm and the periocular mesenchyme, leaving only a small band of Lef1 expression next to the Pax6-expressing RPE (arrowheads).

We next explored the source of Wnt ligands for specifying the proximal RPE by combining *Wnt1-Cr*e and *Le-Cre* to abolish Wnt secretion from both the lens ectoderm and the periocular mesenchyme. Compared to *Wls^ΔLE^* embryos, *Le-Cre*;*Wnt1-Cr*e;*Wntless^flox/flox^* (*Wls^ΔLE+ΔPM^*) mutants exhibited further shortening of the RPE that led to a severe hypopigmentation of the eye (Fig. 6D). This was confirmed by Pax6 and Sox2 staining, which showed extensive invasion of the NR into the backside of the *Wls^ΔLE+ΔPM^* mutant eye cup, leaving only a vestigial segment of the RPE in the center (Fig. 6E, arrowheads). As Wls was ablated in the lens, the surface ectoderm and the periocular mesenchyme, Lef1 expression was also largely eliminated in these regions, except in a small band of mesenchymal cells next to the remaining RPE (Fig. 6E, arrowheads). These results demonstrated that the lens ectoderm, in partial redundancy with the periocular mesenchyme, induces a Wnt signaling gradient in the eye cup to specify the CM and the distal RPE.

### Balanced FGF and Wnt signaling drives CM fate

Our results thus far demonstrated that the eye cup is biased toward the fate of the NR, RPE or CM depending on the availability of FGF and Wnt signaling. What then is the default fate of the eye cup when both FGF and Wnt signaling are absent? To answer this question, we first abolished Wnt signaling by deleting β-catenin in *α-Cre*;*β-catenin^floxlflox^* (*βcat^ΔRet^*) mutants. Consistent with previous studies (*7, 8*), this transformed the CM into the NR as evident by the expansion of Sox2 and loss of Cdo and Msx1 without incurring invasion of Otx2 and Pcad (Fig. 7A). As a result, the pERK level was also elevated in the distal retina. We next ablated both FGF receptors and β-catenin in *α-Cre*; *Fgfr1^flox/flox^;Fgfr2^flox/flox^*;*β-catenin^flox/fox3^* (*Fgfr^ΔRet^*;*βcat^ΔRet^*) embryos. Despite the loss of pERK in the β-catenin negative domain, these *Fgfr^ΔRet^*;*βcat^ΔRet^* mutant cells retained the same NR-like expression patterns as *βcat^ΔRet^* mutants (Fig. 7A), indicating that the eye cup developed into the NR in the absence of FGF and Wnt signaling.

**Figure 7.**
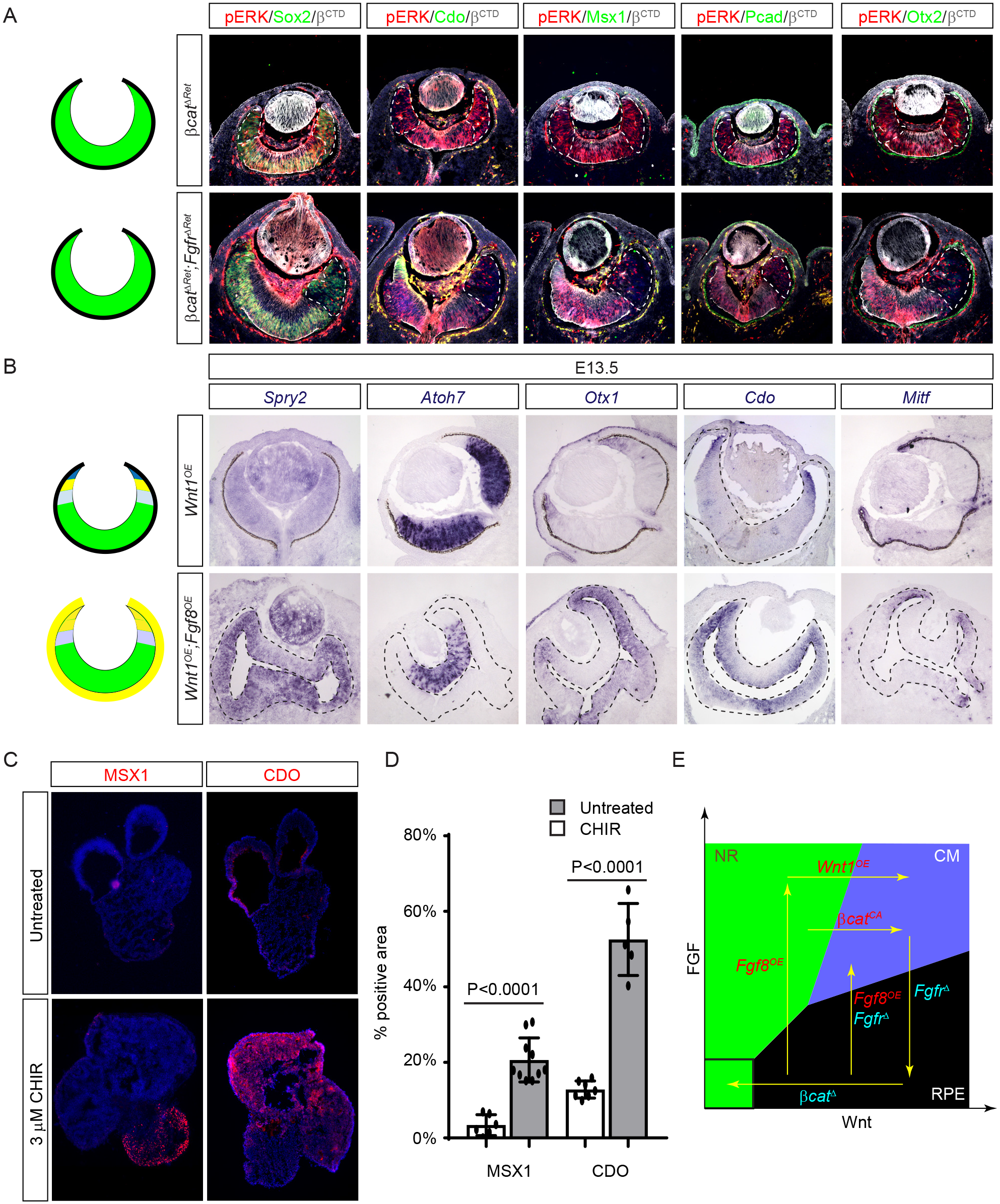
FGF and Wnt induces phase transition to generate the CM. **(A)** Deletion of either β-catenin alone (*βcat^ΔRet^*) or β-catenin and Fgf receptors together (*Fgfr^ΔRet^*;*βcat^ΔRet^*) transformed the distal retina into the NR as indicated by loss of Cdo and Msx1, lack of Otx2 and Pcad and expansion of Sox2. The β-catenin or β-catenin/pERK deficient regions are marked by dotted lines. **(B)** Although overexpression of *Wnt1* in *Wnt1^OE^* embryos did not produce any phenotype in the retina, when combined with overexpression of *Fgf8*, it transformed the RPE domain of the *Wnt1^OE^*;*Fgf8^OE^* eye cup into the CM as indicated by the up regulation of the CM specific genes *Otx1* and *Cdo* and the lack of the RPE marker *Mitf* and the NR marker *Atoh7*. **(C)** The untreated human iPSC organoid culture contained few MSX1^+^ or CDO^+^ CM-like cells. Addition of the 3 µM WNT agonist CHIR99021 significantly expanded the number of MSX1^+^ and CDO^+^ cells. **(D)** Quantification of the MSX1^+^ and CDO^+^ area as the percentage of the total organoid culture. **(E)** FGF and Wnt signaling promote the abrupt and reversible transition of eye cup progenitors into three phases, the NR, the CM and the RPE. Our study showed that the NR can be transformed by constitutive activation of Wnt signaling (*βcat^CA^*) into the CM, which is converted to the RPE after the loss of FGF signaling (*Fgfr^Δ^*). The RPE reverts back to the CM by titrating FGF signaling (*Fgf8^OE^*;*Fgfr^Δ^*). Otherwise, ablation of Wnt signaling (*βcat^Δ^*) or overexpression of *Fgf8* (*Fgf8^OE^*) can turn the RPE into the NR, which may transition further into the CM after overexpression of *Wnt1* (*Wnt1^OE^*).

We next explored the fate of the eye cup when FGF and Wnt signaling are elevated simultaneously. We first bred *α-Cre*;*R26^LSL-Wnt1^* (*Wnt1^OE^*) animals to overexpress *Wnt1* in the retina. Likely because the Wnt transporter Wls is restricted to the distal CM, *Wnt1^OE^* embryos did not exhibit any ocular phenotypes (Fig. 7B), consistent with the above result that the retinal Wnt is dispensable for eye cup patterning (Fig. S4). In contrast, co-expression of both *Wnt1* and *Fgf8* in *α-Cre*;*R26^LSL-Wnt1^*;*R26^LSL-Fgf8^* (*Wnt1^OE^*;*Fgf8^OE^*) mutants led to thickening of the presumptive RPE, which exhibited ectopic FGF signaling activity as indicated by the expression of *Spry2* (Fig. 7B). Unlike *Fgf8^OE^* mutants (Fig. 4C), however, the transformed RPE region in *Wnt1^OE^*;*Fgf8^OE^* mutants failed to express the NR marker *Atoh7*, but up regulated the pan CM markers *Otx1* and *Cdo* at the expense of the RPE marker *Mitf* (Fig. 7B). As reducing the dosage of ectopic Fgf8 signaling can also transform the RPE into the CM, these results demonstrated that the CM fate is specified by the relative, not the absolute, levels of FGF and Wnt signaling.

With this understanding of FGF and Wnt function in CM development, we sought to improve Sasai’s induction-reversal method to generate CM-like stem cells from human embryonic stem cells (hESCs) (*17*). Instead of sequential induction of RPE and NR tissues by adding and then removing GSK and FGFR inhibitors, we hypothesized that, in the absence of FGFR inhibitor, exogenous WNT agonist may cooperate with endogenous FGF produced by the retinal organoid to directly promote the CM fate (*32*). Using human induced pluripotent stem cells (hiPSCs), we confirmed that the standard retinoid differentiation protocol without GSK and FGFR inhibitors produced only a small amount of CM-like tissue as indicated by the limited MSX1 and CDO expression (Fig. 7C). Addition of WNT agonist CHIR99021 (3 µM) drastically increased the number of MSX1 and CDO positive cells in the organoid culture (Fig. 7C and D). Therefore, WNT signaling in the presence of FGF promotes CM differentiation in human retinal organoids.

## DISCUSSION

In this study, we have shown that both FGF and Wnt are transformative signals in eye-cup patterning, forming the basis of a combinatorial code that determines the ocular cell fate. This can be depicted in a two-dimensional map on the strength of FGF and Wnt signaling (Fig. 7E). As demonstrated by our genetic analysis, stimulation of Wnt signaling by constitutively active β-catenin transforms the NR into the CM, which can be further converted to the RPE after removal of FGF receptors. The RPE can revert back to the NR by deleting β-catenin or overexpressing FGF. Otherwise, FGF overexpression may be coopted to transform the RPE to the CM by either the overexpression of Wnt1 or the ablation of FGF receptors, which either balances or reduces the strength of FGF signaling, respectively. Thus cell fate determination at the eye cup stage is a reversible process dictated by the balance between FGF and Wnt signaling, but it is also highly tolerant of extrinsic noise as long as these signals stay within certain confines. Such a switch-like behavior resembles the principle of phase transition in physics, in which matter is maintained in discrete states of solid, liquid or gas within broad ranges of temperature and pressure. Once the boundary of these domains is crossed, however, these physical matters abruptly transition into another phase in an all-or-none manner. We propose that eye cup patterning also acts in a mode of phase transition, a mechanism that may be broadly applicable to other biological systems to achieve versatile yet stable outcomes. As demonstrated by our success in inducing hiPSCs into the CM fate, systematic exploration of biological phase maps may provide insightful guidance for tissue engineering.

In vertebrates from teleost fish to avian, the CM not only contributes to the NR and the peripheral ocular structures during embryogenesis, but also maintain stem cells in adulthood that could replenish retinal neurons after injury (*33*). The regenerative capacity of the CM appears to be lost in adult mammals, but recent studies suggested that, at least in embryonic mice, the CM could give rise to both neuronal and non-neuronal cells (*34, 35*). Previous scRNAseq analyses of mouse retina have readily identified RPCs and neurogenic progenitors that yield retinal neurons, but the CM progenitor pool remains elusive (*20, 21*). By enriching for the peripheral retina, our scRNAseq study has mapped the CM progenitor population, showing that they reside in the same G2/M phase of the cell cycle as neurogenic progenitors. Interestingly, our velocity analysis revealed that neurogenic progenitors inevitably exit cell cycle to differentiate. In contrast, CM progenitors have the bivalent potential to either differentiate or to resume the cell cycle, the latter decision produces daughter cells that may then branch into the neurogenic route. This explains why the CM progenitors can generate both CM cells and retinal neurons, suggesting that these progenitors resemble stem cells in their ability to either self-renew or differentiate. This stem cell-like capacity apparently requires FGF signaling, since loss of FGF signaling will bias CM progenitors toward terminal differentiation. As the regenerative capacity wanes from lower vertebrates to mammals, it will be interesting to explore whether FGF signaling is important for the adult CM to preserve the stem cell-like capability.

Despite their stark differences in anatomical structures, both vertebrate and Drosophila eyes acquired dark pigments to shield them from unwanted light exposure. In Drosophila eye, a Wnt signaling gradient induced by the adjacent head capsule is necessary for emergence of the pigment rim (*3*). In the mouse eye cup, we show that both the lens ectoderm and periocular mesenchyme are sources of Wnt ligands. An earlier study concluded that Wnt ligands from the lens ectoderm regulate proliferation but not differentiation of the optic cup rim (*26*). Our analysis instead reveals that the lens ectoderm-derived Wnt is responsible for specification of the distal RPE and the CM. This is consistent with classic embryological studies that the lens epithelium has the capacity to induce the iris and ciliary body in the optic cup (*36, 37*). Our study thus demonstrates an evolutionarily conserved requirement for paracrine Wnt signaling to induce differentiation of peripheral ocular structures.

In contrast to the conserved role of Wnt signaling, FGF signaling appears to be uniquely required for peripheral eye development in vertebrates. In fact, our study showed that graded FGF signaling is necessary for subdivision of the CM into the proximal, medial and distal zones, which may correspond to pars plana and pars plicata of the ciliary body and the iris in adult eyes. We found that FGF signaling maintains the strength of Wnt signaling, which may serve to extend the range of Wnt signaling gradient. Importantly, FGF signaling stabilizes β-catenin independently of its GSK3β degradation motif, revealing a previously unrecognized mechanism of Wnt signaling regulation (*38*). On the other hand, our study also showed that genetic ablation of both FGF and Wnt signaling transforms the entire retina into the NR. This is reminiscent of NR transformation by deleting β-catenin in the RPE, which lacks FGF signaling (*4, 5*). Thus we propose that the ground state of the eye cup is the NR, which explains why the 3D ES cell culture in the low growth-factor (serum-free) condition naturally develops into the NR (*39*). Only a combination of FGF and Wnt signaling can induce CM formation in the eye cup. These insights are instrumental in our rational design of the CM differentiation protocol for human iPSC culture. Our study shows that FGF and Wnt signaling cooperate to determine the demarcation and subdivision of the eye cup into the NR, CM and RPE. It demonstrates how cross talk between two signaling pathways are able to generate remarkable complexity in organ development.

## MATERIALS AND METHODS

### Mice

All animal procedures were performed according to the protocols approved by the Columbia University’s Institutional Animal Care and Use Committee. *β-catenin^fl3^* was obtained from Dr. Stavroula Kousteni (Columbia University), *Fgf3 ^flox^* from Dr. Suzanne L Mansour (University of Utah), *Fgfr2^flox^* from Dr. David Ornitz (Washington University Medical School), *Mek1^flox/flox^* and *Mek2^-/-^* from Dr. Jean Charron (Universitè Laval), *Pax6 α-Cre (α-Cre)* from Dr. Nadean Brown (Children’s Hospital Research Foundation), *Pax6 Le-Cre (Le-Cre)* from Dr. Richard Lang (Children’s Hospital Research Foundation), *R26^LSL-Fgf8^* from Dr. Yiping Chen (Tulane University), *R26^LSL-Wnt1^* from Dr. Thomas Carroll (UT Southwestern), and *Ai9*, *β-Catenin ^flox^*, *Fgfr1^flox^*, *Msx1-Cre^ERT2^, Wls ^flox^* and *Wnt1-Cre* from Jackson lab (*28, 29, 40-49*). Mice were maintained on a mixed genetic background. At least three animals were analyzed for each of the crosses described. We did not observe phenotypic variations between Cre-heterozygous controls and no-Cre homozygous controls. These two genotypes are hence described together as controls.

### Histology and immunohistochemistry

Histology and immunohistochemistry were performed on the paraffin and cryosections as previously described (*50, 51*). Briefly, slides were washed in PBS 3 times for 5 minutes each, followed by PBST (0.3% Triton) washes 3 times for 5 minutes each before blocked in 10% Horse Serum in PBST (blocking buffer) for 1 hour. Primary antibodies listed in Table 2 were diluted in blocking buffer according to the dilutions and added to the slides in 4C overnight. Antigen retrieval was performed in 10mM Sodium Citrate 0.05% Tween, pH 6.0 in an antigen retriever (Aptum Biologics) for 3 cycles of heating. Secondary antibodies were added the next day for 2 hours at room temperature followed by 5 minutes incubation with DAPI. Slides were cover-slipped using NPG-Glycerol mounting medium for imaging. The antibodies used are listed in Table 2.

**Table 1:**
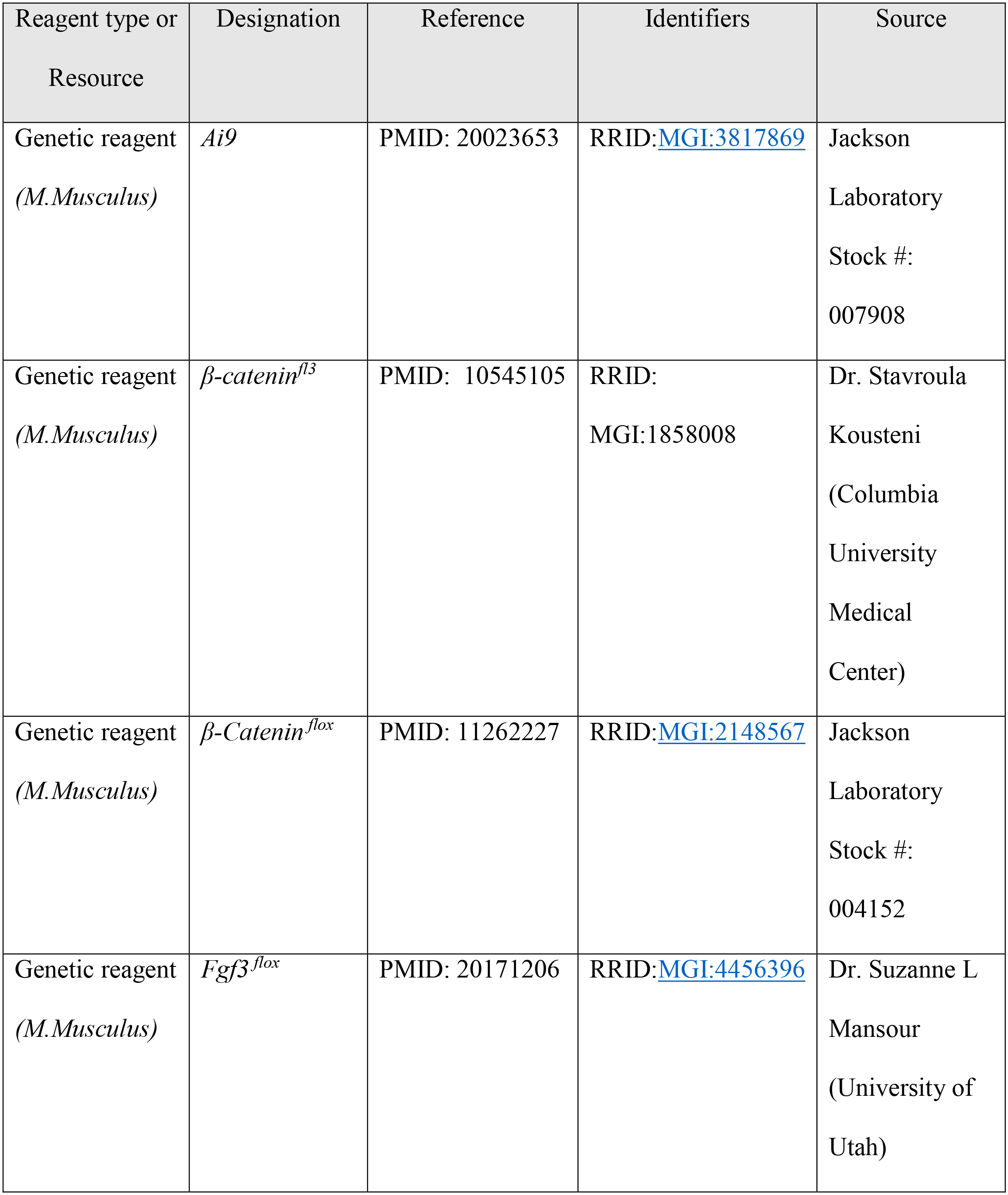
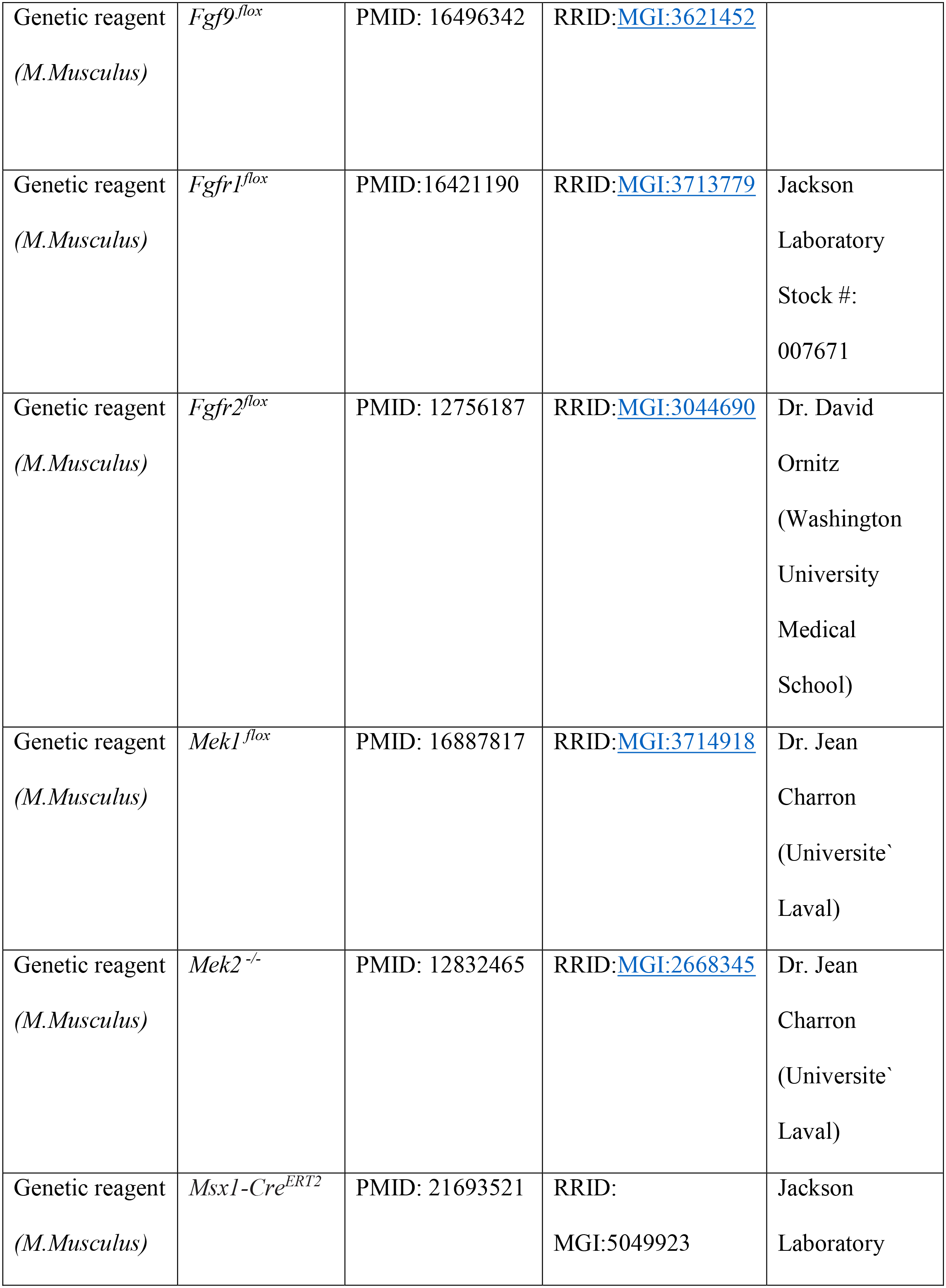
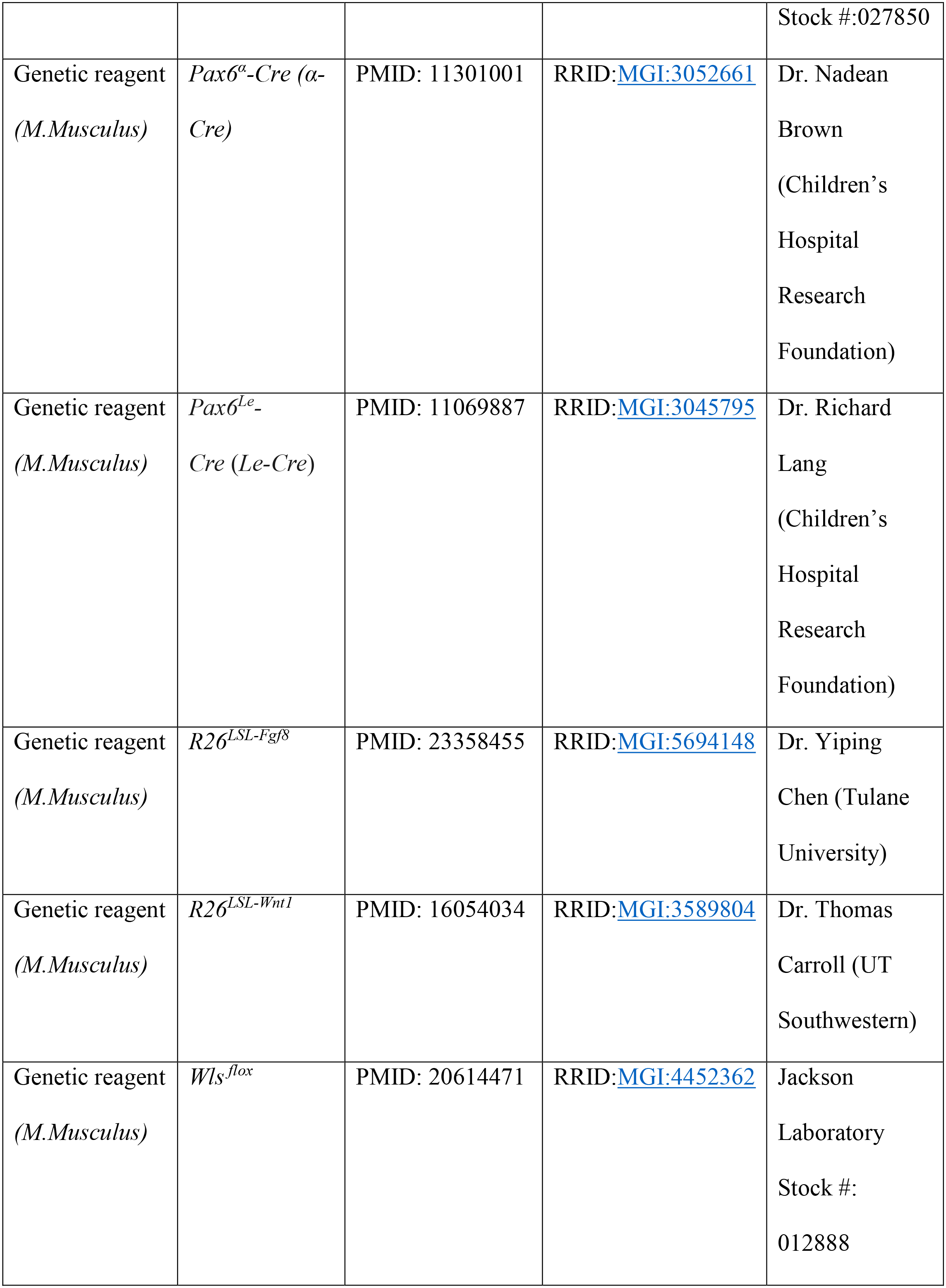
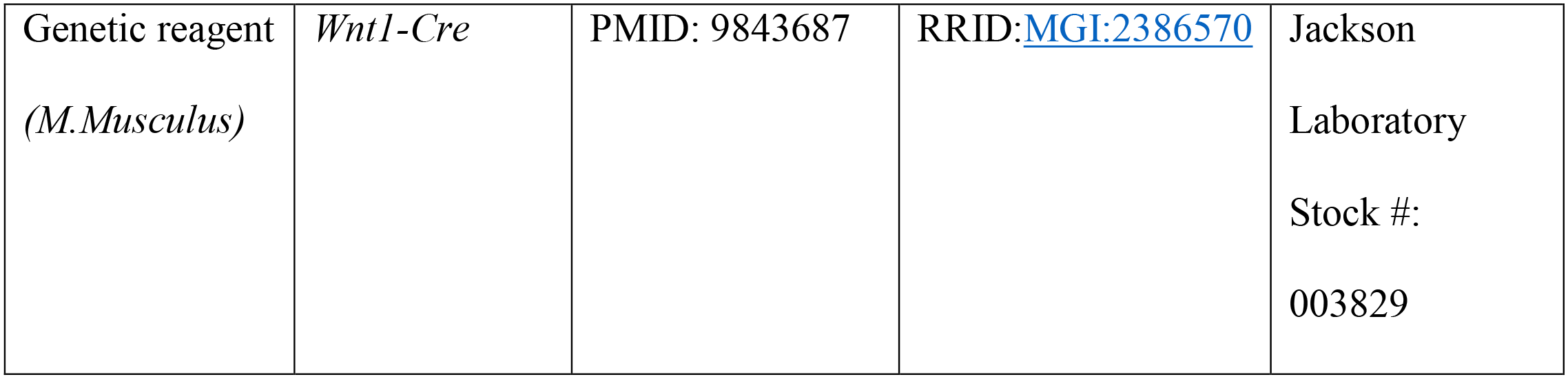
List of Mice Used in the Study.

**Table 2:** List of Antibodies used in the Study.

| Reagent type<br>or Resource | Designation | Source | Identifiers | Additional<br>Information |
| --- | --- | --- | --- | --- |
| Antibody | Goat anti- Cdo | R&D Systems | RRID:AB_2078891 | IHC 1:1000 |
| Antibody | Rabbit anti-Cx43 | Cell Signaling<br>Technology | RRID: AB_2294590 | IHC 1:400 |
| Antibody | Rabbit anti-β-<br>catenin ($\beta^{\text{CTD}}$) | Sigma-Aldrich | RRID: AB_476831 | IHC 1:200 |
| Antibody | Rabbit anti-Non-phospho<br>β-catenin<br>($\beta^{\text{Ser45}}$) | Cell Signaling<br>Technology | RRID: AB_2650576 | IHC 1:200 |
| Antibody | Mouse anti-BrdU | Developmental<br>Studies<br>Hybridoma Bank | RRID:AB_1157913 | IHC 1:200 |
| Antibody | Rabbit anti-GFP | Thermo Fisher<br>Scientific | RRID:AB_221569 | IHC 1:1000 |
| Antibody | Chick anti-GFP | Aves Labs | RRID:AB_10000240 | IHC 1:1000 |
| Antibody | Rabbit anti-Laminin | Sigma-Aldrich | RRID: AB_477163 | IHC 1:1000 |
| Antibody | Rabbit anti-Lef1 | Cell Signaling Technology | RRID:AB_659971 | IHC 1:200 |
| Antibody | Rabbit anti-Mitf | Abcam | RRID:AB_10902226 | IHC 1:100 |
| Antibody | Goat anti-Msx1 | R&D Systems | RRID:AB_2148804 | IHC 1:100 |
| Antibody | Rabbit anti-NICD | Cell Signaling Technology | RRID: AB_ 2153348 | IHC 1:100 |
| Antibody | Goat anti-Otx2 | Santa Cruz Biotechnology | RRID:AB_2157183 | IHC 1:500 |
| Antibody | Rabbit anti-RFP | Rockland Immunochemicals | RRID: AB_828390 | IHC 1:1000 |
| Antibody | Rabbit anti-Pax6 | Covance Research Products Inc. | RRID:AB_291612 | IHC 1:200 |
| Antibody | Goat anti-Pcad | R&D Systems | RRID: AB_355581 | IHC 1:200 |
| Antibody | Rabbit anti-pERK | Cell Signaling Technology | RRID:AB_2095853 | IHC 1:200 |
| Antibody | Rat anti-Sox2 | Thermo Fisher Scientific | RRID:AB_11219471 | IHC 1:500 |
| Antibody | Rabbit anti-Wls | Seven Hills<br>Bioreagents | Catalogue number:<br>RLAB-177 | IHC 1:200 |

### In-situ hybridization

Cryo-sections were processed for in-situ hybridization using previously established protocols (*52*). Briefly, DIG-labeled mRNA *in situ* probes diluted in hybridization buffer was added to the slides and incubated at 65C overnight, followed by washes of high stringent buffers and MABT before blocking with 10% Goat Serum in MABT. Anti-DIG antibody diluted in MABT (with 1% Goat Serum) was added to the slides for 4C overnight. Slides were then washed in MABT followed by Alkaline Phosphatase buffer (NTMT). BM Purple (Millipore Sigma) was added and the color reaction was allowed to develop overnight. Slides were then briefly washed in PBS and cover-slipped with mounting medium for imaging. The probes used are listed in Table 3.

**Table 3:**
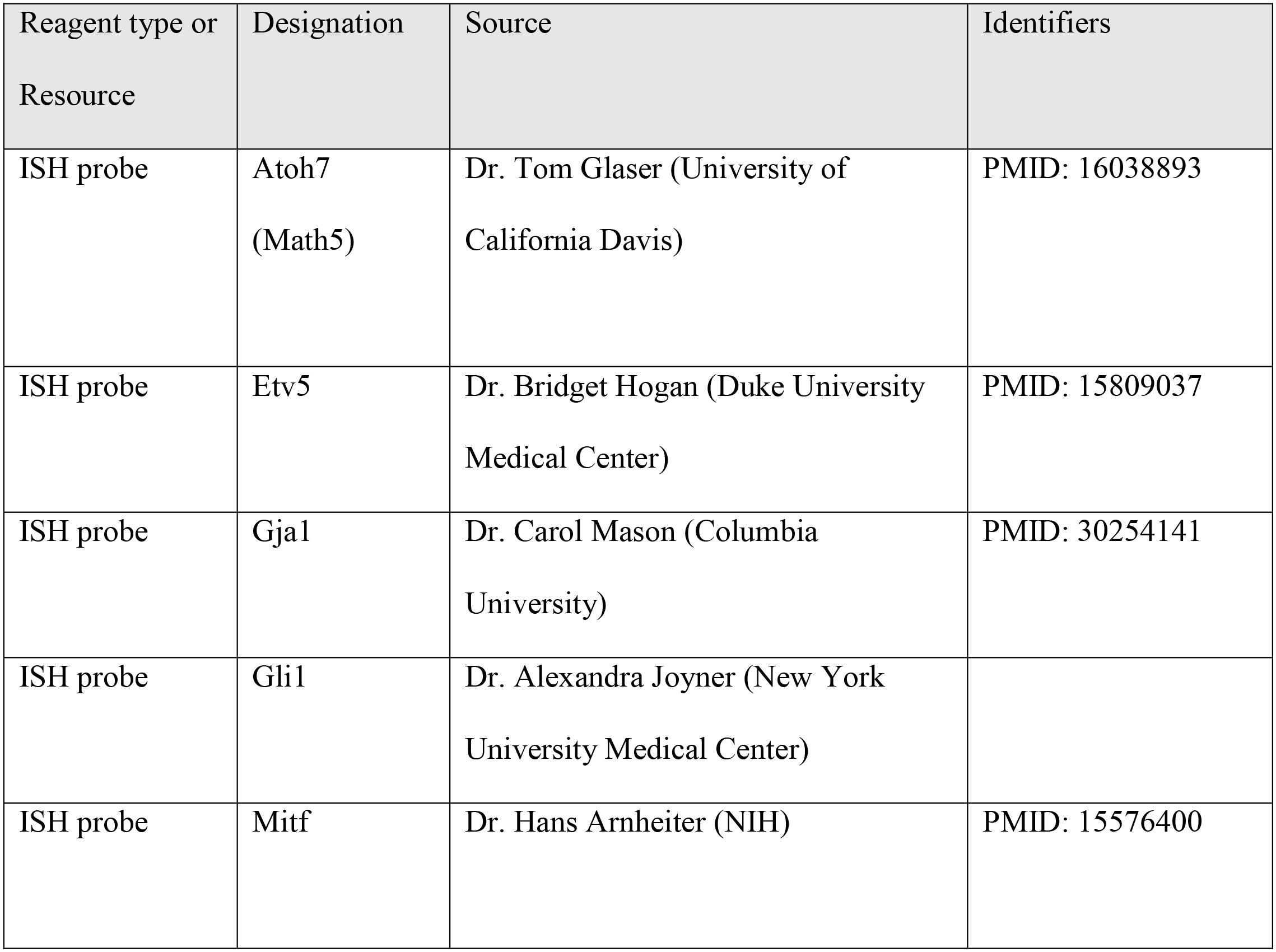
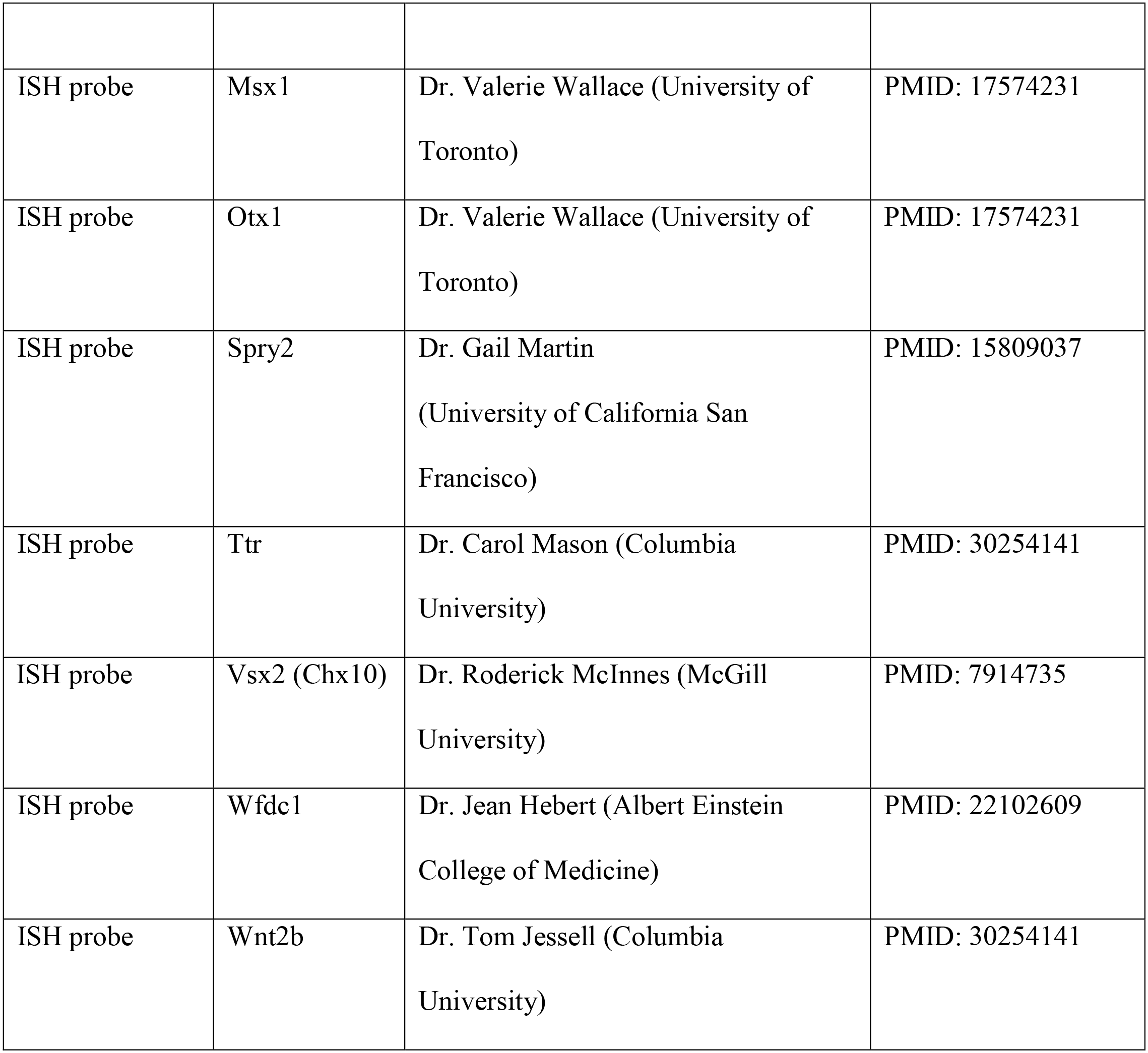
List of ISH probes used in the study.

### Retina dissociation and Flow cytometry

E13.5 retinae with attached RPE from 3 control and mutant embryos each were harvested in ice cold HBSS after removal of the lens. Dissociation was performed using papain dissociation kit (Worthington Biochemical) for 20 minutes at 37°C and stopped using DMEM + 10% FBS. DNase I was added to the samples followed by incubation at 37°C for 5 minutes. Cells were gently triturated using P1000 pipet, centrifuged at 300g and washed with DMEM + 10% FBS. After passing through a 40µm filter and washed again in DMEM + 10% FBS, the cell suspension was with SYTOX to mark dead cells before flow cytometry performed at the Columbia University stem cell core facility. Single cells were collected with Bio-Rad S3e and gated for live cells, GFP positive cells and TdTomato positive cells using FlowJo software. Approximately 100,000 cells were collected into 1.5 ml tubes pre-coated with FBS and containing DMEM + 10%FBS for submission for single cell sequencing.

### Single-cell sequencing

Single-cell sequencing was performed at the single cell sequencing core in the Columbia Genome center. Flow sorted single cells were loaded into Chromium microfluidic chips with v3 chemistry and barcoded with a 10X Chromium controller (10X Genomics). RNA from the barcoded cells was reverse-transcribed and sequencing libraries constructed with Chromium Single Cell v3 reagent kit (10X Genomics). Sequencing was performed on a NovaSeq 6000 (Illumina). The RNAseq data have been deposited at the GEO database (GSE139904).

Raw reads mapped to the mm10 reference genome by 10X Genomics Cell Ranger pipeline (v2.1.1) using default parameters was used for all downstream analyses using Seurat v3 and Velocyto. Briefly, the data set was filtered to contain cells with at-least 200 expressed genes and genes with expression in more than 3 cells. Cells were also filtered for mitochondrial gene expression (<20%). The data-set was log-normalized and scaled. Cells with Td-Tomato reporter expression were extracted and a separate expression matrix was constructed. This provided the ‘control’ and ‘mutant’ gene expression matrices. Unsupervised clustering was performed initially, followed by the manual annotation of Seurat clusters. Biological incompatibility based on gene expression was used to identify doublets. Previously known unique gene expression in extra-retinal cells such as the cornea, lens, and mesenchyme were used to identify extra-retinal clusters. An integration analysis was performed to compare and analyze the ‘control’ and ‘mutant’ gene expression matrices, which included a normalization for cell numbers. The unsupervised clustering with a resolution parameter of 0.7 for both control and mutant cells were represented on a common UMAP space and cluster identity was assigned. Expression of various known genes were used to determine cluster identities. Cell-cycle was assessed using the vignette from Seurat.

For RNA velocity analysis, Velocyto.py was run following an example notebook (https://github.com/velocyto-team/velocyto-notebooks/blob/master/python/ DentateGyrus.ipynb) using the clustering from Seurat. Briefly, loom files were constructed by extracting the reads from the 10X single-cell dataset. Following gene filtering, spliced and unspliced counts were normalized based on total counts per cell. Velocity plots are represented as the transcriptomic integration of control and mutant datasets, to represent similarities and differences in their velocities respectively. Velocity vector fields of control and mutant datasets were calculated by pooling the unspliced and spliced counts from similar cells using the kNN imputation method with 120 neighbors.

### Retinal organoid culture

The hiPSC line PLA-1-3 was maintained on Matrigel (BD)-coated plates in mTeSR medium (STEMCELL Technologies) and passaged with ReleSR (STEMCELL Technologies) (*53*). Retinal organoid differentiation was carried out as previously reported with minor modifications (*54, 55*). In brief, iPSCs at 90% confluence were checkerboard-scraped, approximately 1-1.5 mm^2^, using a 200 µl pipette tip and lifted using a cell scraper. Colony fragments were collected and incubated with (±)blebbistatin in mTeSR medium overnight before moving to Neural Induction Medium 1 (NIM)-1 to form Embryoid Bodies (EBs) in floating culture over three days. All floating culture steps were performed in poly-HEMA (Sigma) coated wells. On differentiation day (DD) 7, floating EBs were transferred to Matrigel-coated wells till DD28, with transition from NIM-1 to NIM-2 medium at DD16. Neuroepithelia were lifted using the checkerboard-scraping method (*56*). From DD28 to DD32, retinal organoids were sorted into poly-HEMA-coated wells and maintained in NIM-2 until the end of the experiment. For ciliary margin induction, differentiation day 32 retinal organoids were treated with GSK inhibitor CHIR 99021 (3 µM) for 10 days of treatment, followed by two days of no treatment before collection. Organoids were washed twice with PBS (Phosphate Buffer saline), fixed for 30 min 4% paraformaldehyde in PBS and cryoprotected by subsequent incubations of 30 min in 15% and 30% sucrose in PBS. The organoids were embedded in Tissue-Tek O.C.T Compound (Sakura Finetek), frozen and stored at −80°C. Sections of 10 µm were generated with a Leica CM1850 cryostat for immunohistochemistry. The CM area was identified by the MSX1^+^ and CDO^+^ expression pattern and measured as the percentage of the total area of organoids in Image J. The statistical significance was calculated using the student’s t test and results are expressed as mean ± s.d. NIM-1: 48.95 mL DMEM/F12 supplemented with 0.5 mL 100x N2 supplement, 0.5 mL 100x Minimum Essential Media-Non Essential Amino Acids (MEM NEAAs), 10 µL 10 mg/mL Heparin (Sigma). NIM-2: 96 mL DMEM/F12 (3:1) supplemented with 2 mL 50x B27 Supplement, 1 mL 100x NEAA, 1 mL 100x Pen Strep (10,000 units/mL of penicillin, 10,000 µg/mL of streptomycin).

## Acknowledgements

The authors thank Drs. Nadean Brown, Thomas Carroll, Jean Charron, Yiping Chen, Stavroula Kousteni, David Ornitz, Suzanne L Mansour for mice, Drs. Hans Arnheiter, Alexandra Joyner, Jean Hebert, Bridget Hogan, Tom Jessell, Tom Glaser, Gail Martin, Carol Mason, Roderick McInnes and Valerie Wallace for in situ probes. We also thank Drs. Carol Mason and Andrew Tomlinson for careful reading of the manuscript. The work was supported by grant from NIH (EY025933 to XZ). The Columbia Ophthalmology Core Facility is supported by NIH Core grant 5P30EY019007 and unrestricted funds from Research to Prevent Blindness (RPB). RB is a recipient of Knights Templar Eye Foundation Career Starter grant. PMJQ is recipient of a Curing Retinal Blindness Foundation grant and Knights Templar Eye Foundation Career Starter grant. BLD is recipient of Capes PhD scholarship. PJF and QLG were supported by funds from the Swiss National Fund (Ambizione grant PZ00P3_174032 to PJF). QW is recipient of Postdoctoral Fellowship from Natural Sciences and Engineering Research Council of Canada. CT is a recipient of Jonas Scholar award. SHT is supported by Jonas Children’s Vision Care. XZ is supported by Jules and Doris Stein Research to Prevent Blindness Professorship.

---

**Figure S1:**
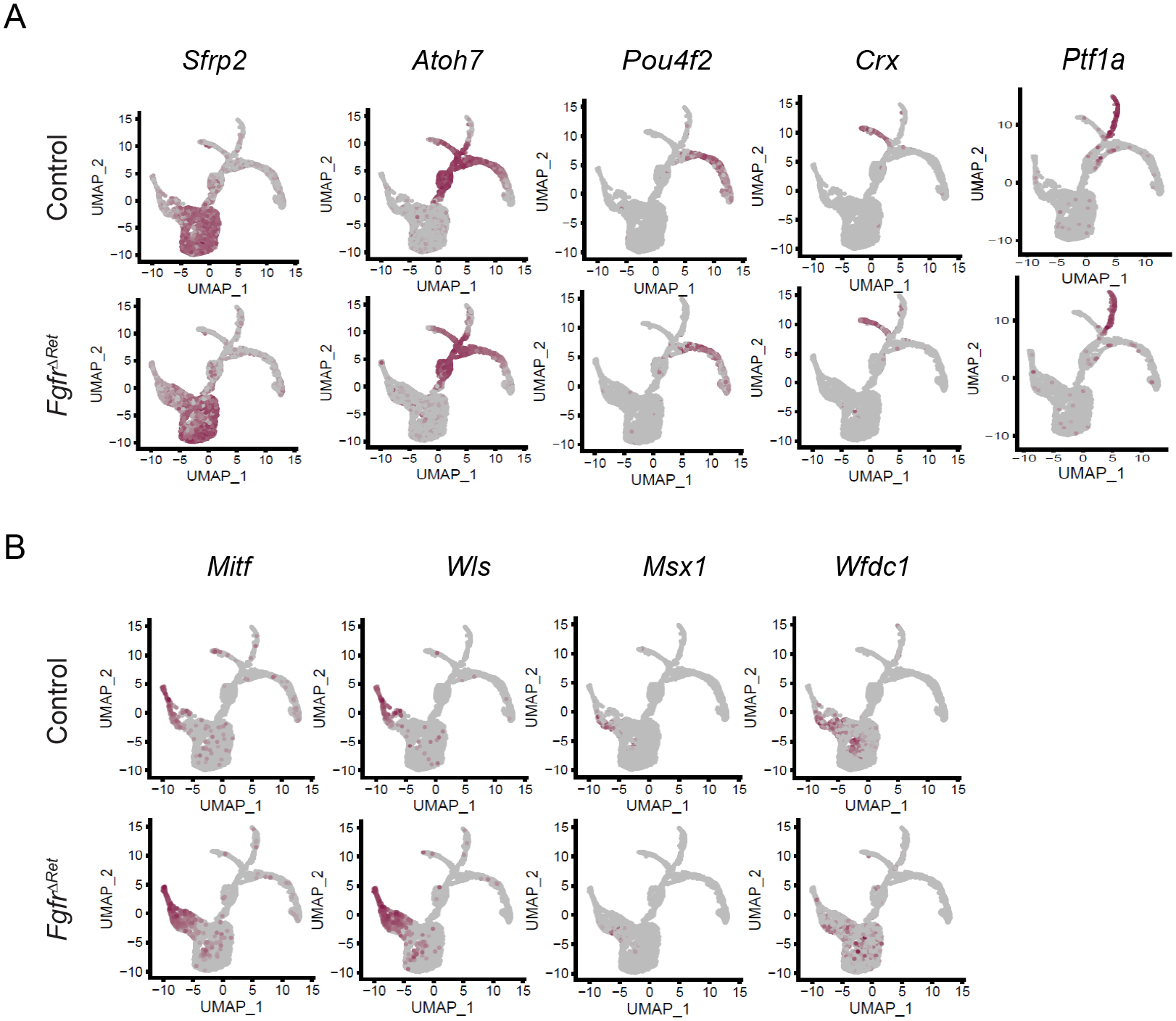
Identification of single-cell clusters by known cell type specific markers. **(A)** Feature plots identified RPC clusters expressing *Sfrp2*, neurogenic clusters expressing *Atoh7*, the RGC cluster expressing *Pou4f2*, the photoreceptor cluster expressing *Crx* and the AC/HC cluster expressing *Ptf1a*. **(B)** Feature plots of the CM specific genes. Notice the increased expression of *Mitf* and *Wls*, the reduced expression of *Wfdc1* and the loss of *Msx1* in *Fgfr^ΔRet^* mutants.

**Figure S2:**
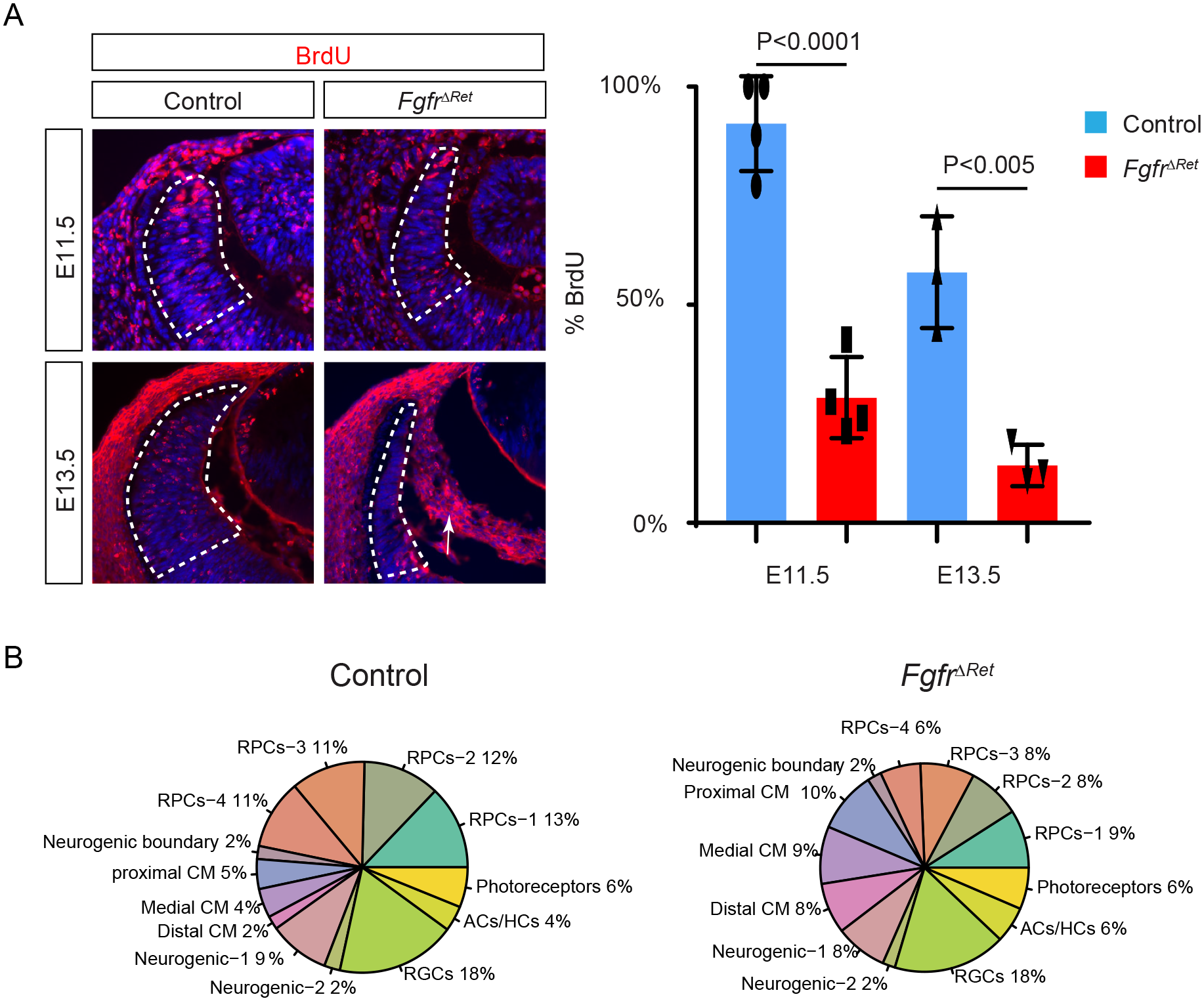
Loss of FGF signaling promoted differentiation of RPCs instead of proliferation. **(A)** The percentages of BrdU+ versus DAPI+ cells were significantly reduced in the peripheral retina (circled in dotted lines) in *Fgfr^ΔRet^* mutants. Arrow points to hyaloid cells in the vitreous. **(B)** The proportions of CM clusters in *Fgfr^ΔRet^* mutants were increased at the expense of RPCs.

**Figure S3:**
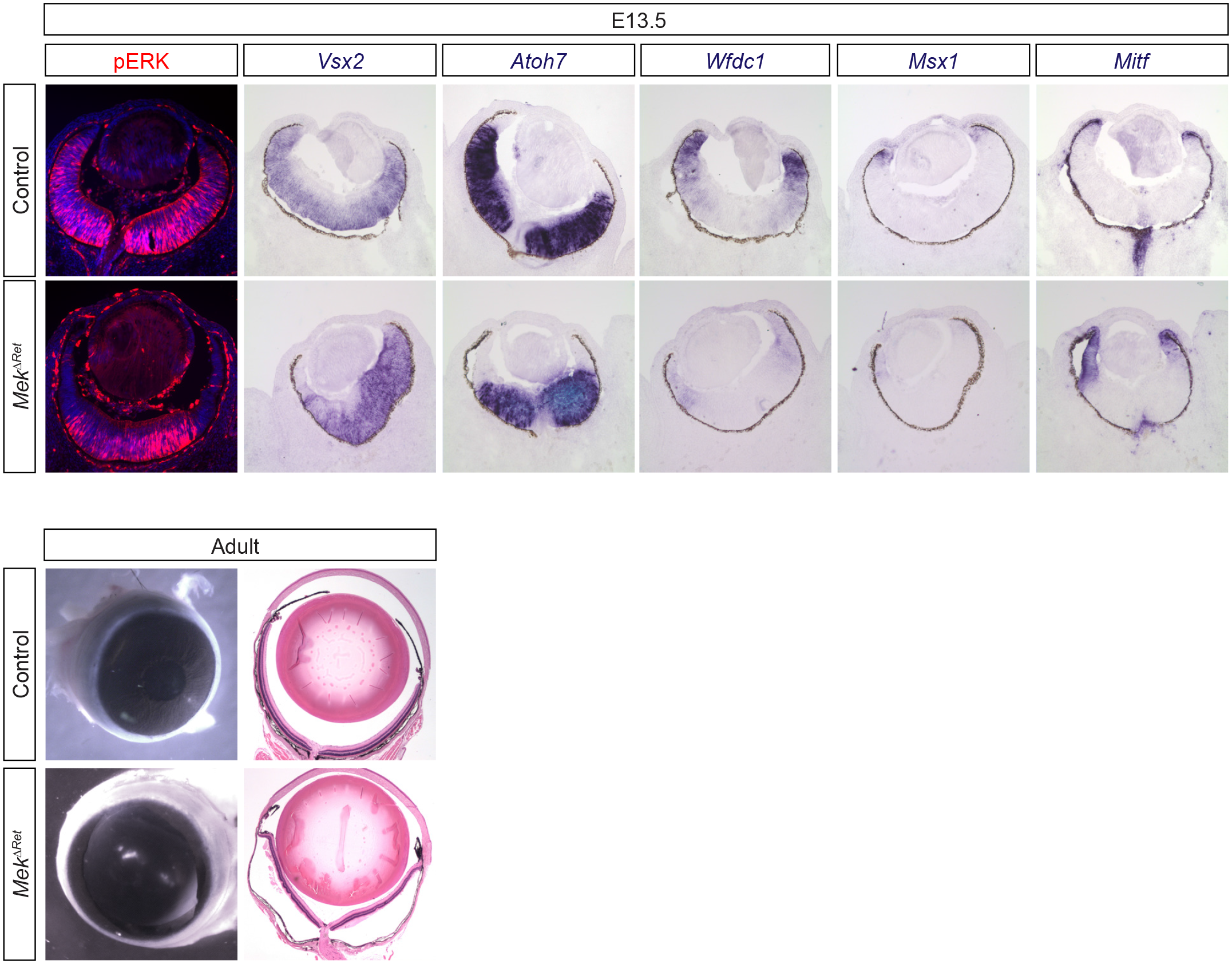
Inactivation of MAPK pathway phenocopied FGF signaling defects. Genetic ablation of *Mek1* and *2* in *Mek^ΔRet^* mutants abolished pERK staining in the peripheral retina. This led to reduced expression of retinal markers *Vsx2* and *Atoh7*, loss of CM markers *Wfdc1* and *Msx1* and expansion of the RPE marker *Mitf*. The adult *Mek^ΔRet^* animals also resemble *Fgfr^ΔRet^* mutants in the iris and ciliary body defects.

**Figure S4:**
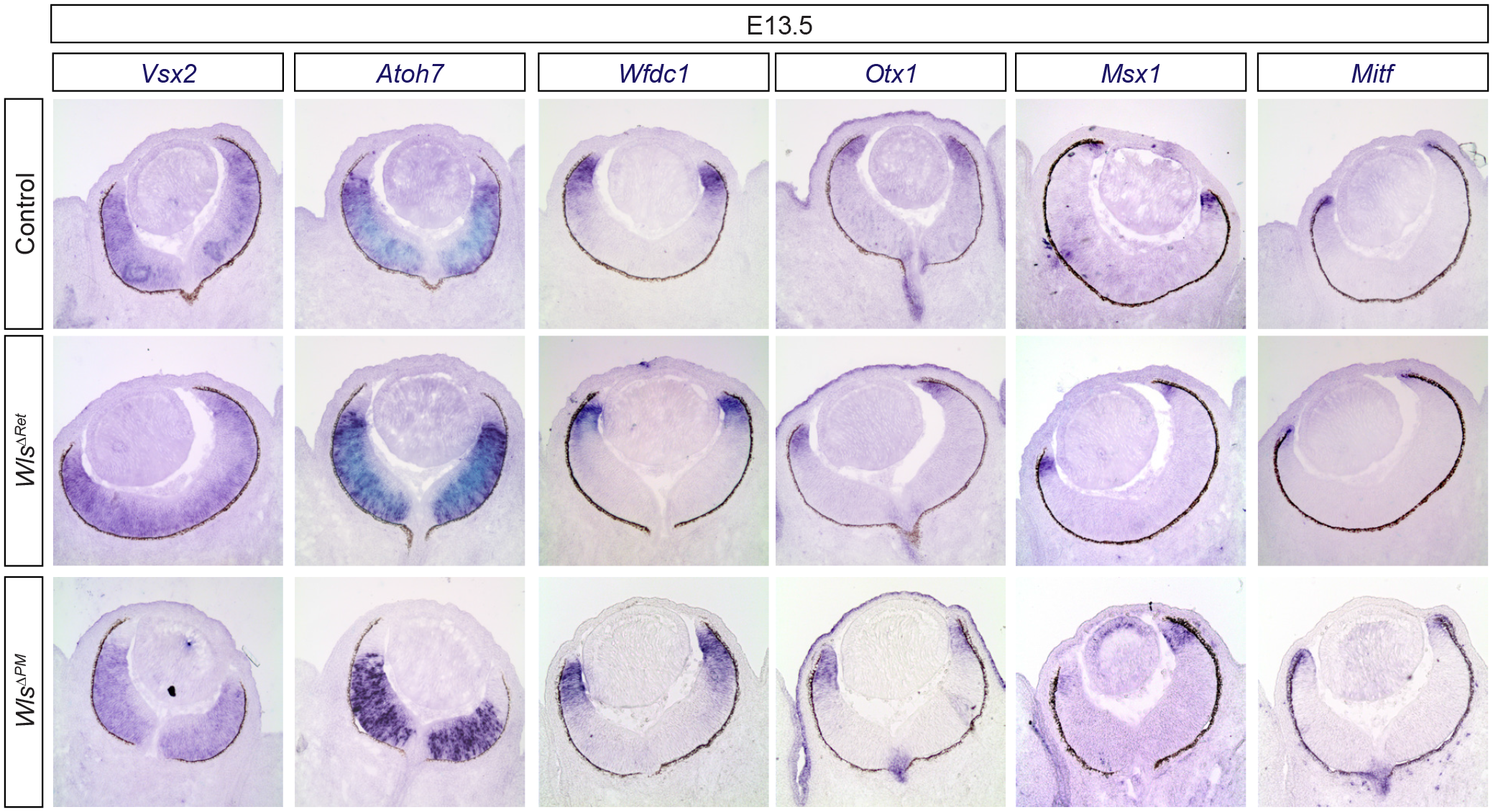
Wls is dispensable in the peripheral retina and the periocular mesenchyme. No retinal phenotype was observed in *Wls^ΔRet^* where Wls was deleted in the peripheral retina and in *Wls^ΔPM^* where Wls was deleted in the periocular mesenchyme.

## REFERENCES

1. R. L. Chow, R. A. Lang, Early eye development in vertebrates. Annu Rev Cell Dev Biol 17, 255–296 (2001).

2. S. Fuhrmann, C. Zou, E. M. Levine, Retinal pigment epithelium development, plasticity, and tissue homeostasis. Exp Eye Res 123, 141–150 (2014).

3. A. Tomlinson, Patterning the peripheral retina of the fly: decoding a gradient. Dev Cell 5, 799–809 (2003).

4. N. Fujimura, M. M. Taketo, M. Mori, V. Korinek, Z. Kozmik, Spatial and temporal regulation of Wnt/beta-catenin signaling is essential for development of the retinal pigment epithelium. Dev Biol 334, 31–45 (2009).

5. P. Westenskow, S. Piccolo, S. Fuhrmann, Beta-catenin controls differentiation of the retinal pigment epithelium in the mouse optic cup by regulating Mitf and Otx2 expression. Development 136, 2505–2510 (2009).

6. K. Bharti et al., A regulatory loop involving PAX6, MITF, and WNT signaling controls retinal pigment epithelium development. PLoS Genet 8, e1002757 (2012).

7. H. Liu et al., Ciliary margin transdifferentiation from neural retina is controlled by canonical Wnt signaling. Dev Biol 308, 54–67 (2007).

8. W. E. Heavner, C. L. Andoniadou, L. H. Pevny, Establishment of the neurogenic boundary of the mouse retina requires cooperation of SOX2 and WNT signaling. Neural Dev 9, 27 (2014).

9. Z. Cai, G. S. Feng, X. Zhang, Temporal requirement of the protein tyrosine phosphatase Shp2 in establishing the neuronal fate in early retinal development. J Neurosci 30, 4110–4119 (2010).

10. Z. Cai et al., Deficient FGF signaling causes optic nerve dysgenesis and ocular coloboma. Development 140, 2711–2723 (2013).

11. S. Chen et al., Defective FGF signaling causes coloboma formation and disrupts retinal neurogenesis. Cell research 23, 254–273 (2013).

12. D. S. Sakaguchi, L. M. Janick, T. A. Reh, Basic fibroblast growth factor (FGF-2) induced transdifferentiation of retinal pigment epithelium: generation of retinal neurons and glia. Dev Dyn 209, 387–398 (1997).

13. C. Pittack, G. B. Grunwald, T. A. Reh, Fibroblast growth factors are necessary for neural retina but not pigmented epithelium differentiation in chick embryos. Development 124, 805–816 (1997).

14. F. Guillemot, C. L. Cepko, Retinal fate and ganglion cell differentiation are potentiated by acidic FGF in an in vitro assay of early retinal development. Development 114, 743–754 (1992).

15. M. Nguyen, H. Arnheiter, Signaling and transcriptional regulation in early mammalian eye development: a link between FGF and MITF. Development 127, 3581–3591 (2000).

16. M. R. Dias da Silva, N. Tiffin, T. Mima, T. Mikawa, J. Hyer, FGF-mediated induction of ciliary body tissue in the chick eye. Dev Biol 304, 272–285 (2007).

17. A. Kuwahara et al., Generation of a ciliary margin-like stem cell niche from self-organizing human retinal tissue. Nat Commun 6, 6286 (2015).

18. J. R. Brewer, P. Mazot, P. Soriano, Genetic insights into the mechanisms of Fgf signaling. Genes Dev 30, 751–771 (2016).

19. T. Marquardt et al., Pax6 is required for the multipotent state of retinal progenitor cells. Cell 105, 43–55 (2001).

20. B. S. Clark et al., Single-Cell RNA-Seq Analysis of Retinal Development Identifies NFI Factors as Regulating Mitotic Exit and Late-Born Cell Specification. Neuron 102, 1111–1126 e1115 (2019).

21. Q. Lo Giudice, M. Leleu, G. La Manno, P. J. Fabre, Single-cell transcriptional logic of cell-fate specification and axon guidance in early-born retinal neurons. Development 146, (2019).

22. G. La Manno et al., RNA velocity of single cells. Nature 560, 494–498 (2018).

23. S. Rowan, C. M. Chen, T. L. Young, D. E. Fisher, C. L. Cepko, Transdifferentiation of the retina into pigmented cells in ocular retardation mice defines a new function of the homeodomain gene Chx10. Development 131, 5139–5152 (2004).

24. S. Zhao et al., Patterning the optic neuroepithelium by FGF signaling and Ras activation. Development 128, 5051–5060 (2001).

25. N. Harada et al., Intestinal polyposis in mice with a dominant stable mutation of the beta-catenin gene. EMBO J 18, 5931–5942 (1999).

26. A. C. Carpenter et al., Wnt ligands from the embryonic surface ectoderm regulate ‘bimetallic strip’ optic cup morphogenesis in mouse. Development 142, 972–982 (2015).

27. A. Ferrer-Vaquer et al., A sensitive and bright single-cell resolution live imaging reporter of Wnt/ss-catenin signaling in the mouse. BMC Dev Biol 10, 121 (2010).

28. R. Ashery-Padan, T. Marquardt, X. Zhou, P. Gruss, Pax6 activity in the lens primordium is required for lens formation and for correct placement of a single retina in the eye. Genes Dev 14, 2701–2711 (2000).

29. P. S. Danielian, D. Muccino, D. H. Rowitch, S. K. Michael, A. P. McMahon, Modification of gene activity in mouse embryos in utero by a tamoxifen-inducible form of Cre recombinase. Curr Biol 8, 1323–1326 (1998).

30. L. Iwai-Takekoshi et al., Activation of Wnt signaling reduces ipsilaterally projecting retinal ganglion cells in pigmented retina. Development 145, (2018).

31. A. N. Smith, L. A. Miller, N. Song, M. M. Taketo, R. A. Lang, The duality of beta-catenin function: a requirement in lens morphogenesis and signaling suppression of lens fate in periocular ectoderm. Dev Biol 285, 477–489 (2005).

32. H. Kinoshita et al., Induction of Functional 3D Ciliary Epithelium-Like Structure From Mouse Induced Pluripotent Stem Cells. Invest Ophthalmol Vis Sci 57, 153–161 (2016).

33. A. J. Fischer, J. L. Bosse, H. M. El-Hodiri, The ciliary marginal zone (CMZ) in development and regeneration of the vertebrate eye. Exp Eye Res 116, 199–204 (2013).

34. M. C. Belanger, B. Robert, M. Cayouette, Msx1-Positive Progenitors in the Retinal Ciliary Margin Give Rise to Both Neural and Non-neural Progenies in Mammals. Dev Cell 40, 137–150 (2017).

35. F. Marcucci et al., The Ciliary Margin Zone of the Mammalian Retina Generates Retinal Ganglion Cells. Cell Rep 17, 3153–3164 (2016).

36. D. C. Beebe, Development of the ciliary body: a brief review. Trans Ophthalmol Soc U K 105 **( Pt** **2**), 123–130 (1986).

37. A. Giroud, [Phenomena of induction & their perturbation in mammals]. Acta Anat (Basel) 30, 297–306 (1957).

38. H. Clevers, R. Nusse, Wnt/beta-catenin signaling and disease. Cell 149, 1192–1205 (2012).

39. M. Eiraku et al., Self-organizing optic-cup morphogenesis in three-dimensional culture. Nature 472, 51–56 (2011).

40. R. V. Hoch, P. Soriano, Context-specific requirements for Fgfr1 signaling through Frs2 and Frs3 during mouse development. Development 133, 663–673 (2006).

41. K. Yu et al., Conditional inactivation of FGF receptor 2 reveals an essential role for FGF signaling in the regulation of osteoblast function and bone growth. Development 130, 3063–3074 (2003).

42. V. Bissonauth, S. Roy, M. Gravel, S. Guillemette, J. Charron, Requirement for Map2k1 (Mek1) in extra-embryonic ectoderm during placentogenesis. Development 133, 3429–3440 (2006).

43. L. F. Belanger et al., Mek2 is dispensable for mouse growth and development. Mol Cell Biol 23, 4778–4787 (2003).

44. Y. Lin, G. Liu, F. Wang, Generation of an Fgf9 conditional null allele. Genesis 44, 150–154 (2006).

45. L. D. Urness, C. N. Paxton, X. Wang, G. C. Schoenwolf, S. L. Mansour, FGF signaling regulates otic placode induction and refinement by controlling both ectodermal target genes and hindbrain Wnt8a. Dev Biol 340, 595–604 (2010).

46. V. Brault et al., Inactivation of the beta-catenin gene by Wnt1-Cre-mediated deletion results in dramatic brain malformation and failure of craniofacial development. Development 128, 1253–1264 (2001).

47. C. Lin et al., Delineating a conserved genetic cassette promoting outgrowth of body appendages. PLoS Genet 9, e1003231 (2013).

48. T. J. Carroll, J. S. Park, S. Hayashi, A. Majumdar, A. P. McMahon, Wnt9b plays a central role in the regulation of mesenchymal to epithelial transitions underlying organogenesis of the mammalian urogenital system. Dev Cell 9, 283–292 (2005).

49. A. C. Carpenter, S. Rao, J. M. Wells, K. Campbell, R. A. Lang, Generation of mice with a conditional null allele for Wntless. Genesis 48, 554–558 (2010).

50. C. Carbe, K. Hertzler-Schaefer, X. Zhang, The functional role of the Meis/Prep-binding elements in Pax6 locus during pancreas and eye development. Dev Biol 363, 320–329 (2012).

51. C. Carbe, X. Zhang, Lens induction requires attenuation of ERK signaling by Nf1. Hum Mol Genet 20, 1315–1323 (2011).

52. C. Carbe et al., An allelic series at the paired box gene 6 (Pax6) locus reveals the functional specificity of Pax genes. J Biol Chem 288, 12130–12141 (2013).

53. Y. Li et al., Patient-specific mutations impair BESTROPHIN1’s essential role in mediating Ca(2+)-dependent Cl(-) currents in human RPE. Elife 6, (2017).

54. P. M. Quinn et al., Human iPSC-Derived Retinas Recapitulate the Fetal CRB1 CRB2 Complex Formation and Demonstrate that Photoreceptors and Muller Glia Are Targets of AAV5. Stem Cell Reports 12, 906–919 (2019).

55. X. Zhong et al., Generation of three-dimensional retinal tissue with functional photoreceptors from human iPSCs. Nat Commun 5, 4047 (2014).

56. C. S. Cowan et al., Cell Types of the Human Retina and Its Organoids at Single-Cell Resolution. Cell 182, 1623–1640 e1634 (2020).

